# Hunger- and thirst-sensing neurons modulate a neuroendocrine network to coordinate sugar and water ingestion

**DOI:** 10.1101/2023.04.06.535891

**Authors:** Amanda J. González-Segarra, Gina Pontes, Nicholas Jourjine, Alexander Del Toro, Kristin Scott

## Abstract

Consumption of food and water is tightly regulated by the nervous system to maintain internal nutrient homeostasis. Although generally considered independently, interactions between hunger and thirst drives are important to coordinate competing needs. In *Drosophila*, four neurons called the Interoceptive Subesophageal zone Neurons (ISNs) respond to intrinsic hunger and thirst signals to oppositely regulate sucrose and water ingestion. Here, we investigate the neural circuit downstream of the ISNs to examine how ingestion is regulated based on internal needs. Utilizing the recently available fly brain connectome, we find that the ISNs synapse with a novel cell type Bilateral T-shaped neuron (BiT) that projects to neuroendocrine centers. *In vivo* neural manipulations revealed that BiT oppositely regulates sugar and water ingestion. Neuroendocrine cells downstream of ISNs include several peptide-releasing and peptide-sensing neurons, including insulin producing cells (IPC), crustacean cardioactive peptide (CCAP) neurons, and CCHamide-2 receptor isoform RA (CCHa2R-RA) neurons. These neurons contribute differentially to ingestion of sugar and water, with IPCs and CCAP neurons oppositely regulating sugar and water ingestion, and CCHa2R-RA neurons modulating only water ingestion. Thus, the decision to consume sugar or water occurs via regulation of a broad peptidergic network that integrates internal signals of nutritional state to generate nutrient-specific ingestion.

## INTRODUCTION

The survival of an organism depends on its ability to coordinate nutrient ingestion with internal nutrient abundance in order to meet its metabolic needs. The nervous system acts as an internal nutrient abundance sensor to drive ingestion in nutrient-deprived states and inhibit ingestion in nutrient-replete states to restore homeostasis (Gizowski & Bourque, 2018, Jourjine, 2017, Sternson, 2013, Qi et al., 2021, Yoshinari et al., 2021). Although generally considered independently, recent studies have demonstrated that interactions between hunger and thirst signals coordinate competing needs (Burnett et al., 2018, Cannell et al., 2016, Jourjine, Mullaney et al., 2016, Watts & Boyle, 2010, Zimmerman et al., 2016).

In mammals, regulation of hunger and thirst drives likely occurs through interactions between food and water ingestion circuits (Eiselt et al., 2021). In the arcuate nucleus of the hypothalamus, neurons that express the agouti related peptide (AgRP) and neuropeptide Y promote food ingestion while neurons that express pro-opiomelanocortin inhibit food ingestion (Aponte et al., 2011, Graham et al., 1997, Sternson, 2013). These neurons can detect circulating ghrelin, glucose, insulin, and leptin secreted from peripheral organs, in addition to receiving input from the gut through the vagus nerve (Sternson, 2013). In the subfornical organ, neurons expressing neuronal nitric oxide synthase (nNOS) promote water ingestion while neurons expressing the vesicular GABA transporter inhibit water ingestion. These cells directly detect blood osmolality and receive input from the gut via the vagus nerve and from the mouth via the trigeminal nerve (Gizowski & Bourque, 2018, Zhang et al., 2022). Interestingly, activation of AgRP neurons decreases water ingestion and inhibition of nNOS expressing cells increases food ingestion (Burnett et al., 2016; Zimmerman et al., 2016). This suggests that hunger sensing cells promote food ingestion and inhibit water ingestion, while thirst sensing cells do the opposite (Jourjine, 2017). However, the underlying circuit mechanisms that lead to this reciprocal coordination of hunger and thirst remain unexplored.

Because of its numerically less complex nervous system, complete connectome, and abundant genetic tools, *Drosophila* is an ideal organism in which to study the coordination of hunger and thirst (Pfeiffer et al., 2008). Like mammals, *Drosophila melanogaster* selectively consumes food when hungry and water when thirsty (Dethier, 1976, Gáliková et al., 2018, Landayan et al., 2021, Lin et al., 2014, Min et al., 2016, Yapici et al., 2016). Moreover, in *Drosophila*, two pairs of neurons, the Interoceptive Subesophageal zone Neurons (ISNs), directly integrate hunger and thirst signals to oppositely regulate sugar and water ingestion (Jourjine, Mullaney et al., 2016).

The ISNs express the adipokinetic hormone receptor (AKHR), a G-protein coupled receptor which binds to the glucagon-like peptide adipokinetic hormone (AKH), a hormone released from the corpora cardiaca during starvation that signals nutrient deprivation (Orchard, 1987, Gáliková et al., 2015). AKH increases ISN activity to drive sugar ingestion and reduce water ingestion. The ISNs also express the TRPV channel Nanchung, which senses changes in hemolymph osmolality. High hemolymph osmolality, such as that experienced during thirst, decreases ISN activity to promote water ingestion and inhibit sugar ingestion (Jourjine, Mullaney et al., 2016). Thus, the ISNs sense both AKH and hemolymph osmolality, arguing that they balance internal osmolality fluctuations and nutrient need (Jourjine, Mullaney et al., 2016). How the ISNs achieve these effects on ingestion remains unclear.

To investigate how the ISNs transform internal nutrient detection into changes in feeding behaviors, we examined the neural network downstream of the ISNs. Using the fly brain connectome, intersectional genetic approaches, *in vivo* functional imaging, and behavioral assays, we identified a neural circuit downstream of the ISNs that regulates sugar and water ingestion. Our work reveals that the ISNs communicate with the neuroendocrine center of the fly brain and regulate the activity of a large number of neurons that transmit or receive peptidergic signals of nutritive state to bidirectionally regulate sugar and water ingestion.

## RESULTS

### The ISNs are peptidergic neurons that release dILP3

To examine how the ISNs reciprocally regulate sugar and water ingestion, we aimed to identify the neural circuit downstream of the ISNs. We first sought to identify which neurotransmitter the ISNs use to communicate with downstream neurons. We expressed RNAi against enzymes involved in neurotransmitter synthesis, vesicular transporters, and neuropeptides in the ISNs and monitored water ingestion in water Interestingly, in an RNAi screen of 18 common neurotransmitters and neuropeptides, only suppression of *Drosophila* insulin-like peptide 3 (dILP3) in the ISNs altered water ingestion (Figure 1A).

**Figure 1.**
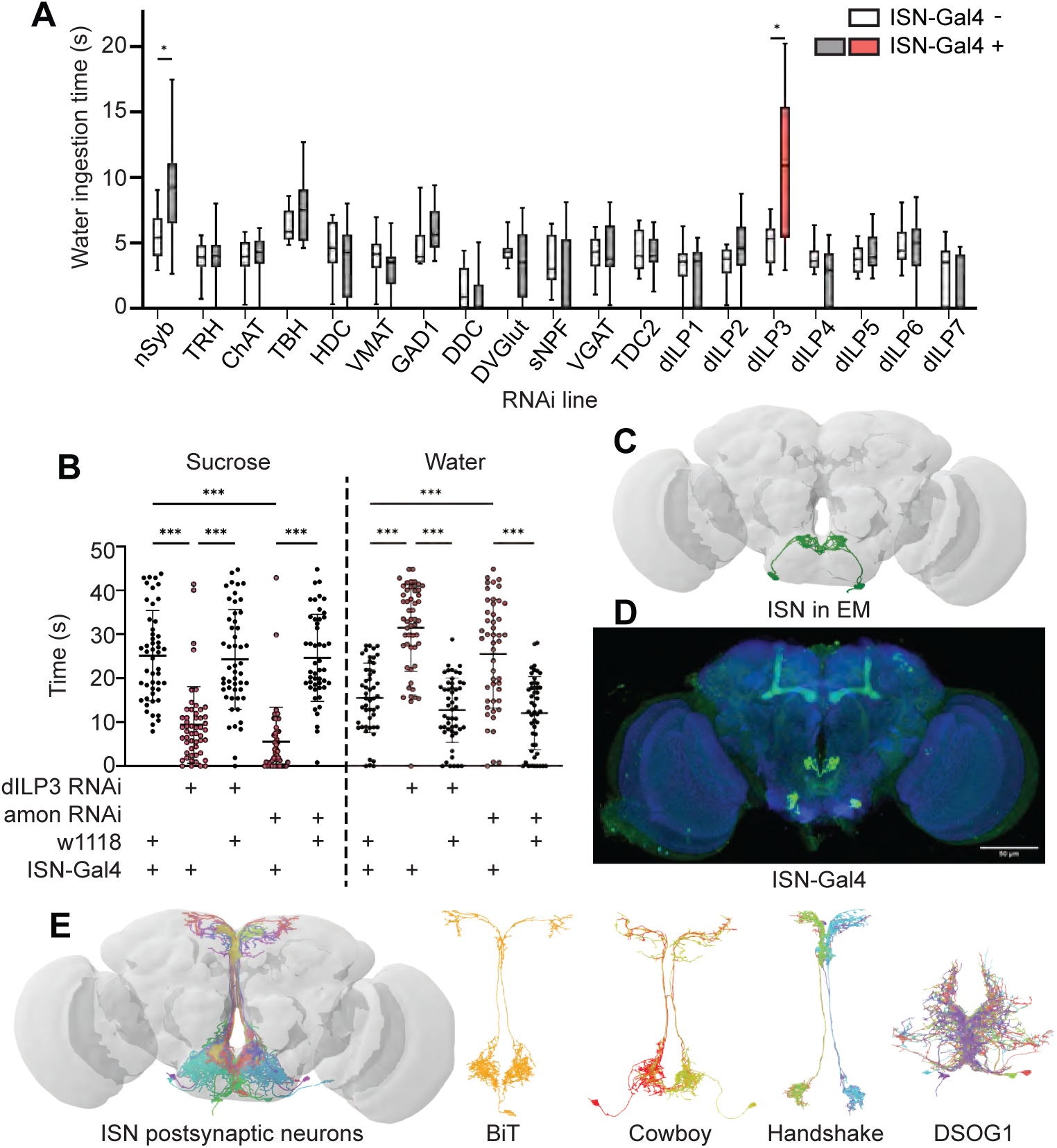
ISNs relay information to the Pars Intercerebralis. (A) Temporal consumption assay screen for water ingestion using RNAi targeting different neurotransmitter pathways. UAS-RNAi + or - ISN-Gal4. RNAi against: nSynaptobrevin (nSyb), tryptophan hydroxylase (TRH), choline acetyltransferase (ChAT), tyrosine beta-hydroxylase (TBH), histamine decarboxylase (HDC), vesicular monoamine transporter (VMAT), glutamic acid decarboxylase 1 (GAD1), dopa decarboxylase (DDC), Drosophila vesicular glutamate transporter (DVGlut), short neuropeptide F (sNPF), vesicular GABA transporter (VGAT), Tyrosine decarboxylase 2 (TDC2), Drosophila insulin like peptide 1 (dILP1), Drosophila insulin like peptide 2 (dILP2), Drosophila insulin like peptide 3 (dILP3), Drosophila insulin like peptide 4 (dILP4), Drosophila insulin like peptide 5 (dILP5), Drosophila insulin like peptide 6 (dILP6), Drosophila insulin like peptide 7 (dILP7). Represented are the mean, and the 10-90 percentile; data was analyzed using Kruskal-Wallis test, followed by multiple comparisons against the RNAi control; p-values were adjusted using False Discovery Rate. n=8-39 animals/genotype except nSyb positive control (70-72). (B) Temporal consumption assay for 1M sucrose or water using RNAi targeting dILP3 or amontillado in ISNs. Sucrose assay: Kruskal-Wallis test followed by Dunn’s multiple comparison tests against ISN control and respective RNAi control. Water assay: ANOVA, Šídák’s multiple comparison test to ISN control and respective RNAi control. n=48-52 animals/genotype. (C) ISNs reconstruction from FAFB volume. (D) Light microscopy image of ISN-Gal4 registered to JFRC2010. (E) ISN postsynaptic neurons based on synapse predictions using FAFB volume (Zheng et al. 2018) and connectome annotation versioning engine (CAVE, Buhmann et al. 2021, Heinrich et al. 2018). Left: 10 postsynaptic neurons, right: postsynaptic neurons BiT, Cowboy, Handshake and DSOG1. *p<0.05, ***p<0.001 deprived flies (Figure 1A). As decreasing activity of the ISNs increases water ingestion (Jourjine, Mullaney et al., 2016), we anticipated that an RNAi against the ISN neurotransmitter would decrease neurotransmission and increase water ingestion.

To confirm that dILP3 functions in the ISNs and to test whether it is involved in the reciprocal regulation of water and sugar ingestion, we expressed RNAi against dILP3 in the ISNs and measured sugar or water ingestion in water sated or thirsty flies respectively (Figure 1B). As an additional approach to reduce dILP3, we expressed an RNAi against a neuropeptide processing protease, *amontillado* (Siekhaus & Fuller, 1999), in the ISNs and tested sugar and water ingestion. We found that knockdown of either dILP3 or *amontillado* in the ISNs caused both a decrease in sugar ingestion and an increase in water ingestion (Figure 1B). This is the same phenotype that was previously reported in the ISNs upon loss of neurotransmission (Jourjine, Mullaney et al., 2016). These data argue that the ISNs are peptidergic neurons that release dILP3 and that one function of dILP3 is to promote sugar ingestion and inhibit water ingestion.

### The ISNs synapse onto neurons that arborize in neuroendocrine and feeding centers

*Drosophila* has one insulin-like receptor (dInR), a tyrosine kinase type receptor homologous to the human insulin receptor, which binds dILP3 and six of the additional *Drosophila* insulin-like peptides (Brogiolo et al., 2001, Claeys et al., 2002, Clancy et al., 2001, Fernandez, et al., 1995, Grönke et al., 2010, Nässel & Broeck, 2016, Tatar et al., 2001). In adult flies, insulin signaling has been shown to regulate an array of physiological processes including metabolism, feeding, reproduction, and lifespan (Badisco et al., 2013, Biglou et al., 2021, Clancy et al., 2001, Nässel et al., 2013, Ohhara et al., 2018). Since dInR is ubiquitous and involved in many different processes (Chen et al., 1996, Garofalo, 2002, Veenstra et al., 2008), we could not leverage neurotransmitter receptor identity for postsynaptic neuron identification. We instead used the *trans*-Tango system (Talay et al., 2017), a genetic trans-synaptic tracer, to label neurons postsynaptic to the ISNs (Supp 1A). We expressed the *trans*-Tango ligand in the ISNs and its receptor panneurally. Binding of the ligand to its receptor induces GFP expression in the receptor-expressing cells and labels potential synaptic partners (Talay et al., 2017). *trans*-Tango labeling revealed numerous ISN postsynaptic arborizations in the subesophageal zone (SEZ), a brain region associated with taste processing and feeding circuits (Gordon & Scott, 2009, Scott et al., 2001, Wang et al., 2004), and along the median bundle to the superior medial protocerebrum (SMP), a neuroendocrine center (Hartenstein, 2006, Nässel & Zandawala, 2020) (Supp 1A). However, as many ISN candidate postsynaptic neurons were labeled, the morphology of individual neurons was unclear.

To comprehensively examine the postsynaptic partners of the ISNs, we employed the Full Adult Fly Brain (FAFB) volume, a whole-brain electron microscopy volume that provides synaptic resolution of all neurons in the fly brain (Zheng, Lauritzen et al., 2018). We manually reconstructed the ISNs using CATMAID (Li et al., 2019, Saalfeld et al., 2009) by tracing neuronal arbors from the pharyngeal nerve with large cell bodies in the SEZ. Due to the ISNs’ unique morphology, with large cell bodies near the pharyngeal nerve and dense neurites in the flange that cross the midline, we used visual morphological comparison of the reconstructed ISNs in the FAFB volume (Figure 1C) and light microscopy images of *ISN-Gal4* (Figure 1D) to identify the ISNs. Once we had reconstructed the ISNs, we labeled presynaptic sites in the ISNs and postsynaptic sites in other neurons based on known synapse active zone structure (Zhai & Bellen, 2004). We then reconstructed neurons that were postsynaptic to the ISNs.

Soon after we had reconstructed the four ISNs and several postsynaptic neurons in CATMAID, the FlyWire whole brain connectome of more than 80,000 reconstructed EM neurons became available (Dorkenwald et al., 2022, flywire.ai). Since FlyWire uses the FAFB volume, we used the coordinates of the ISNs we traced in CATMAID to locate them in FlyWire. Additionally, we compared a pointcloud generated from a registered light microscopy image of *ISN-Gal4* (Figure 1D) to the reconstructed ISNs in the FAFB volume (Figure 1C) to further confirm ISN identity. We identified neurons downstream of the ISNs (Fig 1E). We found that the ISNs have 104 predicted postsynaptic partners with 5 or more synapses, comprising 9 morphological cell types (Supplementary Table 1, Supp 1C-K). These include known cell types (Cowboy, DSOG1, FLAa2, FLAa3/Lgr3 and the ISNs; Lee et al., 2020, Pool et al., 2014, Sterne et al., 2021, Yu et al., 2013) as well as many uncharacterized cell types. The ISN predicted postsynaptic partners include projection neurons that project along the median bundle to the SMP (64 cells), local SEZ neurons (18 cells), ascending neurons with projections coming through the neck connective (10 cells), descending neurons with projections leaving through the neck connective (8 cells), and the ISNs themselves (4 cells). This connectivity is consistent with the connectivity determined by *trans*-Tango (Supp 1A). Overall, the ISN synaptic connectivity suggests that the hunger and thirst signals sensed by the ISNs are conveyed to a broad network, with the potential to coordinate feeding behaviors (SEZ neurons), nutrient status (SMP neuroendocrine centers), and movement or digestion (ascending and descending neurons). We note that as neuropeptides may diffuse over long distances (van den Pol, 2012), ISN dILP3 release may also influence activity of additional neurons that are not synaptically connected to the ISNs.

### The ISN postsynaptic neuron BiT reciprocally regulates sugar and water ingestion

As the majority of the ISN predicted postsynaptic partners project to the SMP, we examined whether ISN communication to this region regulates neuroendocrine cells and/or influences feeding behavior. As a first step, we focused on an uncharacterized neuron that receives the most synaptic input from the ISN per single cell. We named this neuron Bilateral T-shaped neuron (BiT). BiT has its cell body in the SEZ and bilateral projections in the flange and SMP. It receives 7.4% of ISN synaptic output (301/4050 synapses) (Supp 1B and Supplementary Table 1). In turn, the ISNs are the main synaptic input to BiT, comprising 17% of BiT’s synaptic input (301/1763 synapses). We generated a split-Gal4 line that labels BiT to study its function (Fig 2B). We screened over 20 AD-DBD combinations and found that *VT002073-Gal4.AD* and *VT040568-Gal4.DBD* specifically labeled BiT. We confirmed this by comparing a pointcloud generated from a registered light microscopy image of *BiT split-Gal4* (Fig 2B) with the reconstructed BiT in the FAFB volume (Fig 2A).

**Figure 2.**
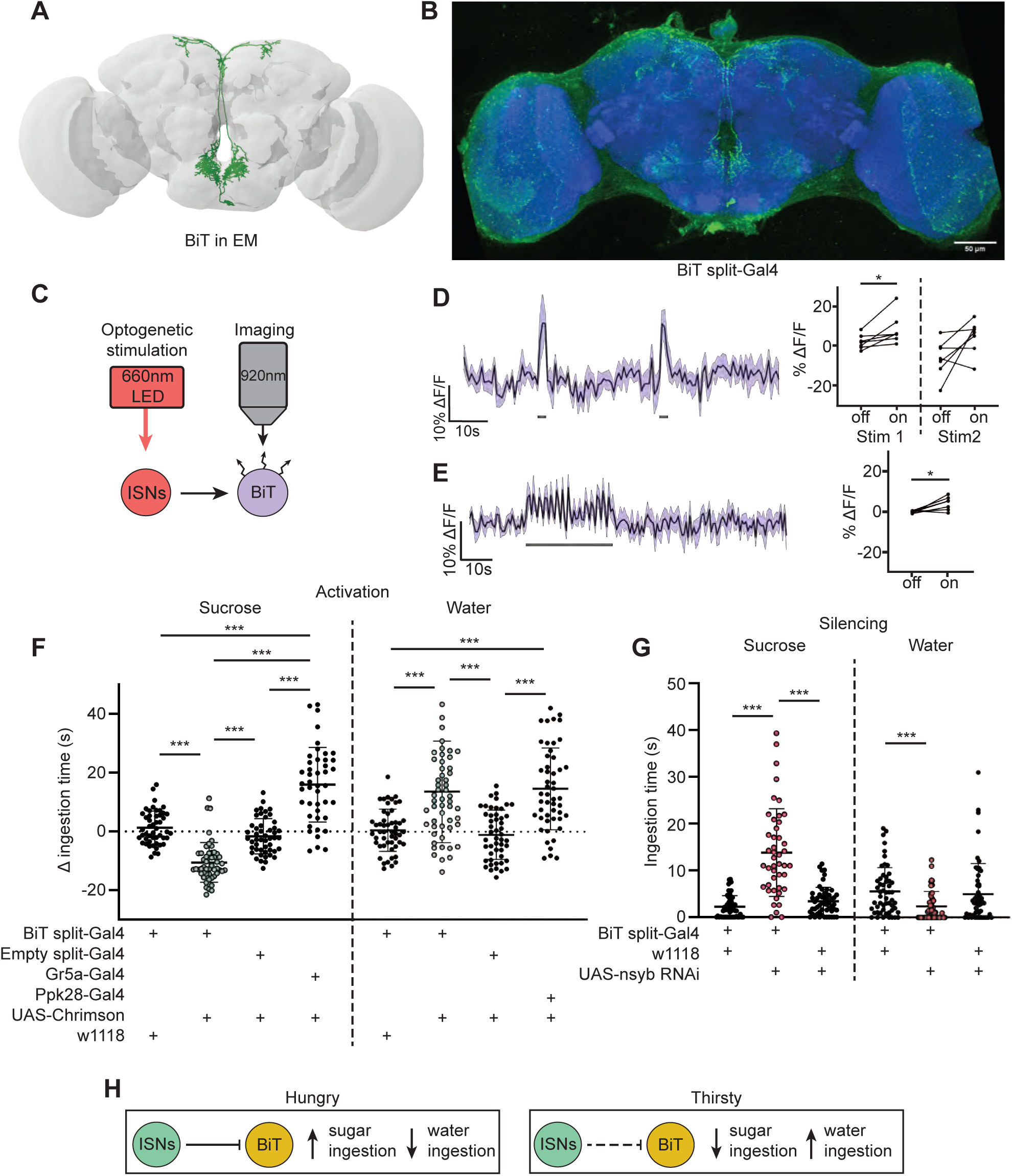
ISNs inhibit BiT, which oppositely regulates sugar and water ingestion. (A) BiT neuron reconstruction from FAFB dataset. (B) Light microscopy image of BiT split-Gal4. (C) Experimental setup for in vivo voltage imaging. We expressed the light sensitive ion channel Chrimson in the ISNs and optogenetically stimulated them with 660nm LED. We expressed the voltage sensor ArcLight in BiT and imaged it with a 2 photon microscope. (D) ArcLight response of BiT soma to 2s optogenetic stimulation of the ISNs or (E) 30s optogenetic stimulation of the ISNs. Left: Scatter plot shows mean +/- SEM of all flies imaged, gray bars represent LED stimulation. Right: Quantification of mean fluorescence intensity before stim (off) and during stim (on), each dot represents one fly. Paired Wilcoxon and paired t-test (Stim 2, p=0.07). n=7 flies. (F) Temporal consumption assay for 1M sucrose or water during acute optogenetic activation of BiT with Chrimson. Ingestion time of females exposed to light normalized to dark controls of indicated genotype. Sucrose: Kruskal-Wallis test with Dunn’s multiple comparison test. Water: One-way ANOVA with Holm-Šídák multiple comparison test. n=44-54 animals/genotype.(G) Temporal consumption assay for 1M sucrose or water using RNAi targeting nSyb in BiT. Kruskal-Wallis with Dunn’s multiple comparison test. n=45-57 animals/genotype. (H) Neural model for BiT coordination of sucrose and water intake. Dashed lines indicate inactive synapses. *p<0.05, ***p<0.001

To test whether the ISNs are functionally connected to BiT, we conducted *in vivo* functional imaging experiments in which we activated the ISNs while simultaneously monitoring BiT’s neural activity. We expressed the light activated cation channel Chrimson in the ISNs and the voltage sensor ArcLight in BiT (Fig 2C) (Jin et al., 2012, Klapoetke et al., 2014). In one experiment, we applied two consecutive 2s stimulations (Fig 2D) to test whether the response was reproducible. In another experiment, we applied a longer 30s stimulation (Fig 2E) to ensure we captured the full response to ISN stimulation since dILPs can act over longer time scales (Sudhakar et al., 2020). In both experiments, we found that stimulating the ISNs increased ArcLight fluorescence in BiT, demonstrating that BiT became hyperpolarized (Fig 2D-E). Oscillation in BiT’s response during the 30s stimulation (Fig 2E) is due to oscillations in the LED stimulation paradigm. Thus, increased activity in the ISNs inhibits BiT.

Next, we tested whether BiT modulates sugar or water ingestion. We measured total ingestion time of sugar or water while activating or inhibiting BiT. We found that acute optogenetic activation of BiT decreased sugar ingestion and increased water ingestion (Fig 2F). Moreover, reducing synaptic transmission in BiT using nSynaptobrevin (nSyb) RNAi caused increased sugar ingestion and decreased water ingestion (Fig 2G). These data demonstrate that BiT is both necessary and sufficient to regulate sugar and water ingestion. Furthermore, we find that the activation and silencing phenotypes for BiT are opposite to the ISN phenotypes, consistent with our calcium imaging studies that the ISNs inhibit BiT. These findings reveal that the coordination of sugar and water ingestion is maintained downstream of the ISNs.

These studies demonstrate that BiT activity reciprocally regulates sugar and water ingestion, similar to the ISNs. Hunger signals (i.e. adipokinetic hormone) activate the ISNs, causing the ISNs to inhibit BiT, which in turn increases sugar ingestion. On the other hand, thirst signals (i.e. high hemolymph osmolality) inhibit the ISNs, releasing ISN inhibition onto BiT, causing an increase in water ingestion (Fig 2H). Strikingly, although BiT is only one ISN downstream neuron, its activity increases and decreases are sufficient to coordinate both sugar and water ingestion, suggesting that it is a critical node in the ISN network.

### BiT downstream partners include neuroendocrine cells that convey nutritional status

To examine how BiT coordinates sugar and water ingestion, we investigated the neural circuit downstream of BiT using the FlyWire connectome (Fig 3). The FAFB connectivity revealed that BiT has 93 predicted postsynaptic partners. Unlike the ISNs’ downstream partners, which only innervate the SMP and SEZ, BiT postsynaptic partners reach more brain regions including the superior lateral protocerebrum (SLP), fan shaped body (FB), lobula, SMP, and SEZ. This suggests that the hunger and thirst signals detected by the ISNs are conveyed by BiT to widely regulate brain activity.

**Figure 3.**
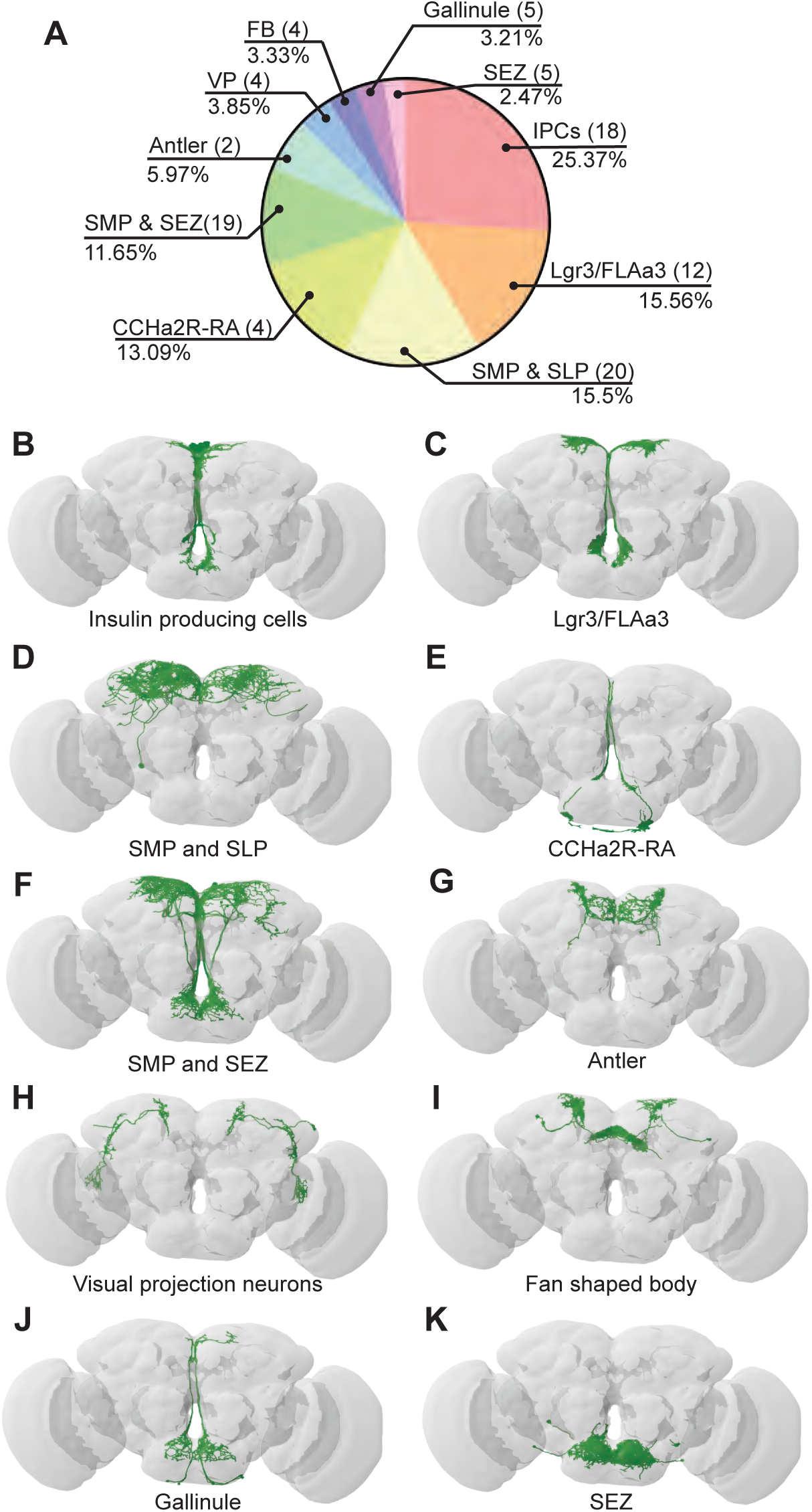
BiT postsynaptic neurons include neuroendocrine cells. (A) Distribution of synaptic output from BiT divided by cell class or brain region. Total of 1742 synapses from BiT and 93 postsynaptic partners. IPCs (18 neurons) receive 25.37% of all BiT output, Lgr3/FLAa3 (12 neurons) 15.56%, SMP and SLP (20 neurons) 15.5%, CCHa2R-RA (4 neurons) 13.09%, SMP and SEZ (19 neurons) 11.65%, Antler (2 neurons) 5.97%, visual projections (4 neurons) 3.85%, fan shaped body (4 neurons) 3.33%, Gallinule (5 neurons) 3.21%, SEZ (5 neurons) 2.47%. Only post-synaptic partners with 5 or more synapses were considered for this analysis. Reconstruction of IPCs (B), Lgr3/FLAa3 neurons (C), neurons innervating the SMP and SLP (D), CCHa2R-RA neurons (E), neurons innervating the SMP and SEZ (F), Antler neurons (G),visual projection neurons (H), neurons innervating the fan shaped body (I), Gallinule neurons (J), and neurons innervating the SEZ (K).

Many of the BiT predicted postsynaptic partners arborize in both the SEZ and SMP, suggesting that they might coordinate nutritional status and feeding. Several BiT targets transmit or receive peptidergic signals of nutrient state. For example, BiT postsynaptic partners include insulin producing cells (IPCs), FLAa3/Lgr3 neurons, and neurons labeled by the *CCHa2R-RA-Gal4* line (Deng et al., 2019) (Fig 3, Supplementary Table 2). IPCs are a well-studied cell type that release dILP2, dILP3 and dILP5, regulate glucose uptake, and influence many physiological processes including feeding (Nässel et al., 2013, Ohhara et al., 2018). FLAa3/Lgr3 neurons detect dILP8 and influence sugar ingestion (Meissner et al., 2016, Yeom et al., 2021, Yu et al., 2013). CCHa2 and its receptor CCHa2R have been shown to participate in feeding regulation and regulate insulin signaling, although the function of CCHa2R-RA neurons has not been examined (Deng et al., 2019, Ida et al., 2012, Ren et al., 2015, Sano et al., 2014, Shahid et al., 2021). Thus, BiT is predicted to synapse onto many neuroendocrine neurons, possibly enabling integration of the hunger and thirst signals sensed by ISNs with diverse nutrient state signals.

### IPCs regulate sugar and water ingestion

The IPCs integrate multiple signals of nutrient status and regulate feeding and metabolism (Nässel & Zandawala, 2020). We found that the ISNs are connected to the IPCs via BiT. BiT is the main synaptic input into IPCs, making up 25% of the IPCs’ synaptic input (442/1735) and IPCs receive 25% of BiT’s synaptic output (442/1742) (Fig 3, Table 2). We tested whether BiT is functionally connected to the IPCs by optogenetically stimulating BiT and monitoring activity in IPCs using the calcium sensor GCaMP6s (Chen et al., 2013). We found that BiT inhibits IPCs (Supp 3A-B), consistent with neurotransmitter predictions (Eckstein et al., 2020) that BiT uses glutamate, which can act as an inhibitory neurotransmitter in *Drosophila* (Liu et al., 2013).

To test whether IPCs modulate ingestion of sucrose or water under conditions that reveal ISN behavioral phenotypes, we measured ingestion time of sucrose or water while acutely activating the IPCs. We found that acute activation of IPCs increased sucrose ingestion and decreased water ingestion (Supp 3E). These results are consistent with one study (Sudhakar et al., 2020) but differ from other studies showing that acute IPC activation limits ingestion of sucrose or food (Nässel et al., 2015, Wang et al., 2020). IPCs integrate many signals and release multiple peptides (Sano et al., 2015, Söderberg et al., 2012, Ohhara et al., 2017, Wang et al., 2020), suggesting that differences in these behavioral results may, in part, stem from differences in the current nutritional state sensed by the IPCs. While further experiments are needed to elucidate how IPCs coordinate nutrient state and ingestion under different conditions, our results show that BiT regulates IPC activity and that IPC activity coordinates both sugar and water ingestion.

### CCHa2R-RA neurons regulate water ingestion downstream of BiT

A number of studies indicate that CCHa2 and its receptor CCHa2R promote food intake and appetite in various insects, including blowflies (Ida et al., 2012), aphids (Shahid et al., 2021) and *Drosophila* (Ren et al., 2015). BiT synapses with CCHa2R-RA neurons, four neurons with cell bodies in the SEZ and arbors in the flange and pars intercerebralis (PI) (Fig 4B). BiT is the dominant input onto CCHa2R-RA neurons, comprising 94% of CCHa2R-RA presynaptic sites (171/181 synapses). CCHa2R-RA neurons receive the most output from BiT per single cell comprising 13% of BiT’s output (228/1742 synapses). To investigate whether BiT’s synaptic input to CCHa2R-RA neurons regulates ingestion, we examined the functional connectivity between BiT and CCHa2R-RA neurons and the behavioral phenotypes associated with CCHa2R-RA neurons.

**Figure 4.**
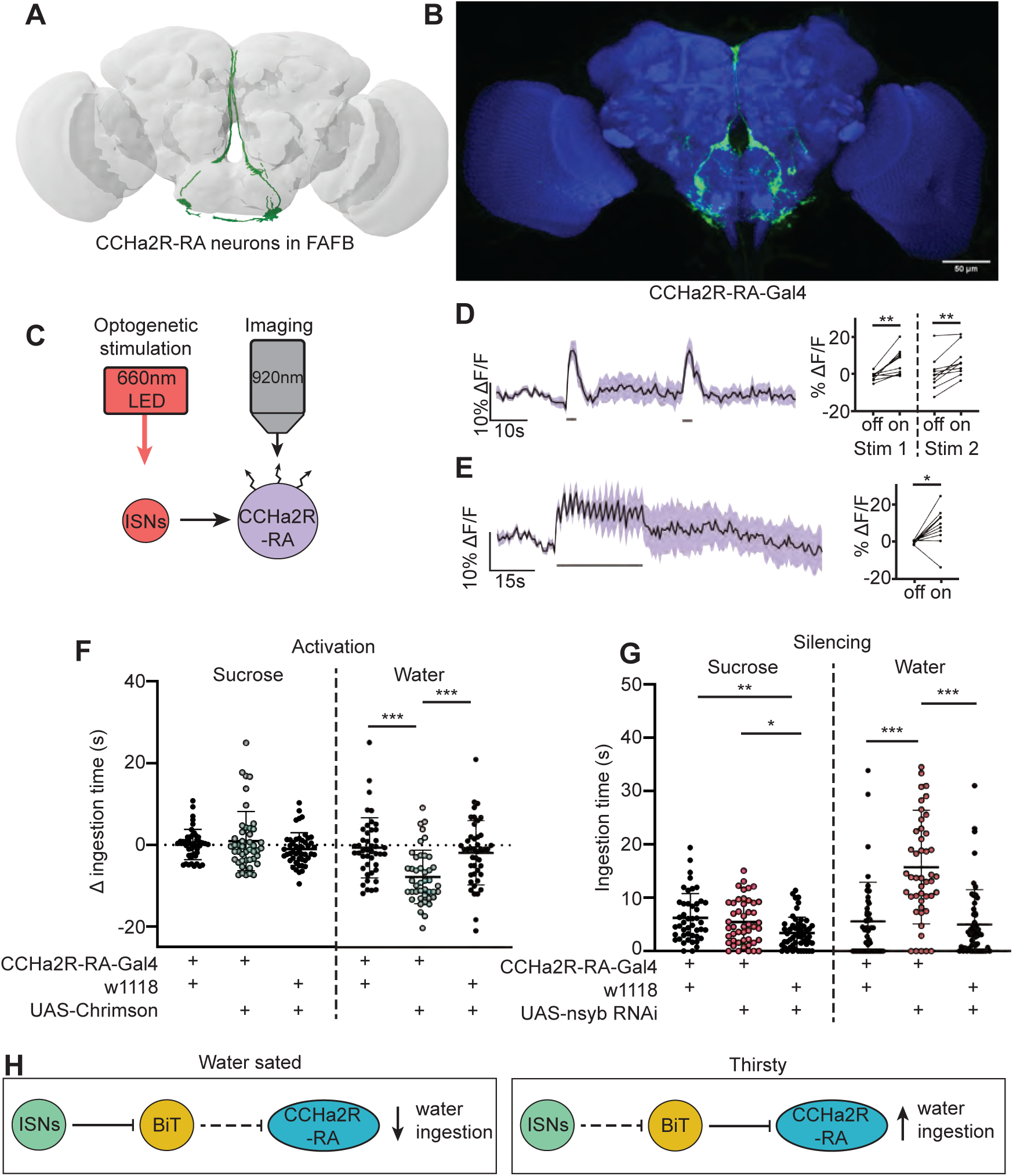
CCHa2R-RA neurons regulate water but not sugar ingestion and are likely inhibited by BiT. (A) CCHa2R-RA neurons reconstruction from FAFB dataset. (B) Light microscopy image of CCHa2R-RA-Gal4. (C) Experimental setup for in vivo calcium imaging. We expressed the light sensitive ion channel Chrimson in the ISNs and optogenetically stimulated them with 660nm LED. We expressed the calcium sensor GCaMP in the CCHa2R-RA neurons and imaged them with a 2 photon microscope. (D) Calcium responses of CCHa2R-RA neurites in SEZ to 2s optogenetic stimulation of the ISNs or (E) 30s optogenetic stimulation of the ISNs. Left: Scatter plot shows mean +/- SEM of all flies imaged, gray bars represent LED stimulation. Right: Quantification of mean fluorescence intensity before stim (off) and during stim (on), each dot represents one fly. Paired t-test and paired Wilcoxon test. n=10 flies. (F) Temporal consumption assay for 1M sucrose or water during acute optogenetic activation of CCHa2R-RA neurons with Chrimson. Ingestion time of females exposed to light normalized to dark controls of indicated genotype. Sucrose: Kruskal-Wallis with Dunn’s multiple comparison test. Water: One-way ANOVA with Holm-Šídák multiple comparison test. n=42-47 animals/genotype. (G) Temporal consumption assay for 1M sucrose or water using RNAi targeting nSyb in CCHa2R-RA neurons. Kruskal-Wallis with Dunn’s multiple comparison test. n=45-54 animals/genotype. (H) Neural model for CCHa2R-RA regulation of water intake. Dashed lines indicate inactive synapses. *p<0.05, **p<0.01, ***p<0.001

We monitored activity in CCHa2R-RA neurons with the calcium indicator GCaMP6s upon optogenetic stimulation of BiT; however, we did not observe a response in CCHa2R-RA neurons (Supp 4D-E). As BiT likely inhibits CCHa2R-RA neurons, it is possible that we were unable to detect an inhibitory response in CCHa2R-RA neurons using a calcium sensor. We therefore monitored activity of CCHa2R-RA neurons upon optogenetic stimulation of the ISNs, as the ISNs should activate CCHa2R-RA neurons given that the ISNs inhibit BiT (Fig 4C). Indeed, we found that CCHa2R-RA neurons showed robust calcium responses upon ISN stimulation (Fig 4D-E), demonstrating that these neurons are functionally connected to the ISNs, likely via BiT inhibition.

To test if CCHa2R-RA neurons regulate sugar or water ingestion, we manipulated activity in these neurons and measured ingestion of sugar or water. We found that activation of CCHa2R-RA neurons decreased water ingestion but did not change sugar ingestion (Fig 4F). Moreover, inhibiting neurotransmission in CCHa2R-RA neurons increased water ingestion (Fig 4G), but did not change sucrose ingestion relative to *CCHa2R-RA-Gal4* controls. These behavioral experiments demonstrate that peptide-sensing neurons downstream of the ISNs regulate water ingestion. The finding that CCHa2-RA neurons recapitulate the water ingestion phenotypes of the ISNs but not sugar ingestion phenotypes suggests that the ISNs activate different arrays of peptidergic neurons that contribute differentially to ingestion of specific nutrients.

### CCAP neurons are downstream of the ISNs and reciprocally regulate sugar and water ingestion

In a separate effort to find neurons that are postsynaptic to the ISNs, we tested whether neurons that had previously been implicated in ingestion were functionally connected to the ISNs. We conducted pilot *in vivo* functional imaging experiments monitoring the activity of candidate neurons with GCaMP7b while optogenetically stimulating the ISNs. We found one set of peptidergic neurons, the crustacean cardioactive peptide (CCAP) neurons, that were activated upon ISN optogenetic stimulation (Fig 5D-E).

**Figure 5.**
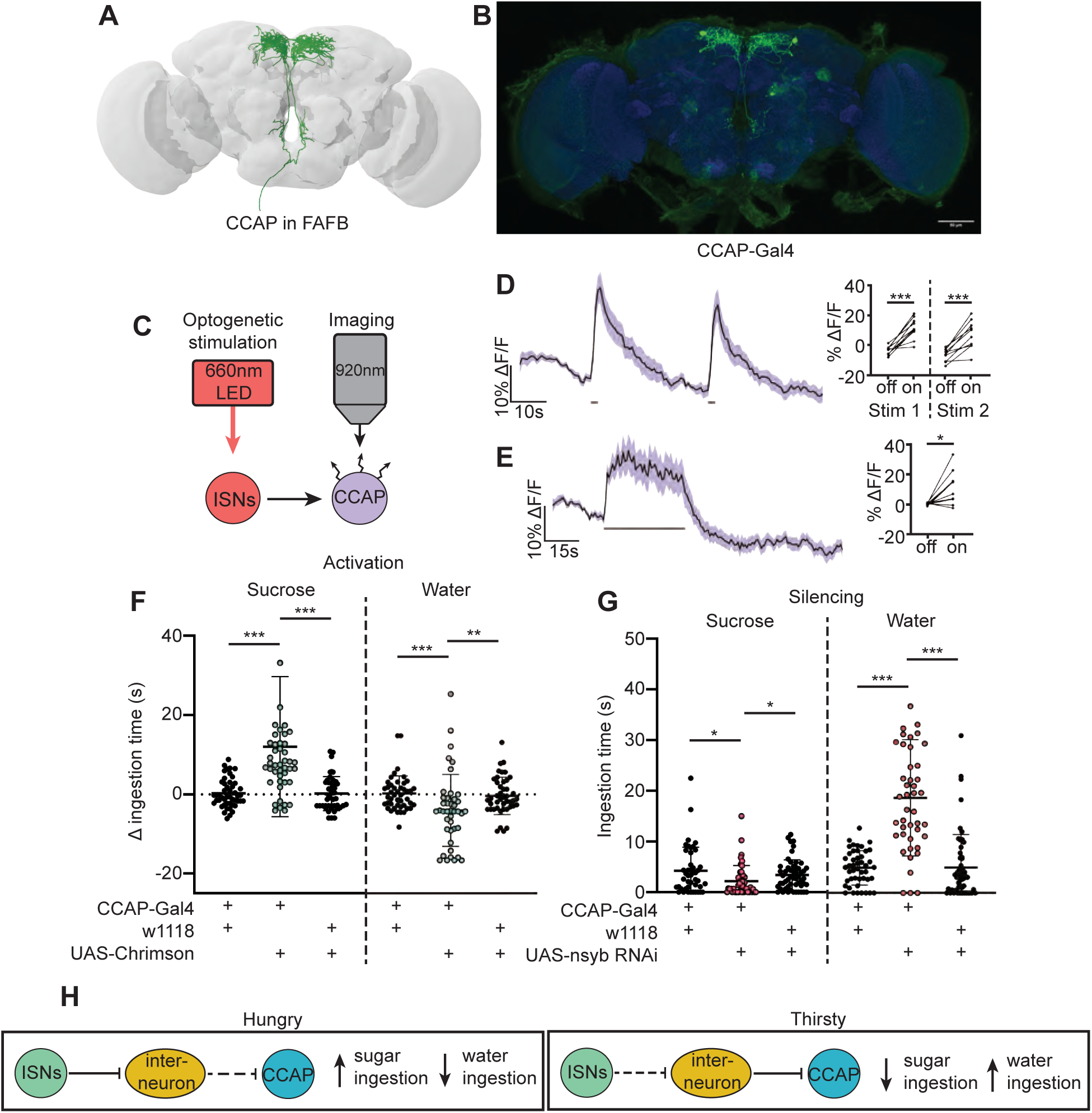
CCAP neurons are downstream of the ISNs and oppositely regulate sugar and water ingestion. (A) CCAP neurons reconstruction from FAFB dataset. (B) Light microscopy image of CCAP-Gal4. (C) Experimental setup for in vivo calcium imaging. We expressed the light sensitive ion channel Chrimson in the ISNs and optogenetically stimulated them with 660nm LED. We expressed the calcium sensor GCaMP in the CCAP neurons and imaged them with a 2 photon microscope. (D) Calcium response of CCAP neurites to 2s optogenetic stimulation of the ISNs or (E) 30s optogenetic stimulation of the ISNs. Left: Scatter plot shows mean +/- SEM of all flies imaged, gray bars represent LED stimulation. Right: Quantification of mean fluorescence intensity before stim (off) and during stim (on), each dot represents one fly. Paired t-test. n=10 flies. (F) Temporal consumption assay for 1M sucrose or water during acute optogenetic activation of CCAP neurons with Chrimson. Ingestion time of females exposed to light normalized to dark controls of indicated genotype. Sucrose: Kruskal-Wallis with Dunn’s multiple comparison test, Water: One-way ANOVA with Holm-Šídák multiple comparison test. n=42-48 animals/genotype. (G) Temporal consumption assay for 1M sucrose or water using RNAi targeting nSyb in CCAP neurons. Kruskal-Wallis with Dunn’s multiple comparison test. n=45-54 animals/genotype. (H) Neural model for CCAP coordination of sugar and water intake. Dashed lines indicate inactive synapses. *p<0.05, **p<0.01, ***p<0.001

CCAP neurons have been shown to regulate feeding behavior in adult *Drosophila* as loss of CCAP in these neurons reduced sucrose ingestion (Williams et al., 2020). To directly test if CCAP neural activity modulates sugar or water ingestion, we acutely manipulated the activity of CCAP neurons and measured ingestion of sugar or water. We found that activation of CCAP decreased water ingestion and increased sugar ingestion (Fig 5F). To test whether CCAP neurons are necessary for sugar and water ingestion, we reduced CCAP neurotransmission with nSyb RNAi, and measured ingestion of sugar or water. We found that silencing CCAP neurons decreased sugar ingestion and increased water ingestion (Fig 5G), demonstrating that CCAP neurons reciprocally regulate sugar and water ingestion, similar to the ISNs.

Although CCAP neurons are functionally connected to the ISNs, their synaptic connectivity is indirect. We identified the CCAP neurons in the FAFB volume (Fig 5A) and found weak connections between CCAP neurons and ISN synaptic partners: Cowboy (5 synapses), VESa1 (22 synapses), and a novel neuron we named Bilateral T-shaped neuron 2, based on its anatomical similarities to BiT (37 synapses). In addition, the ISN third-order neuron CCHa2R-RA neurons provide 26 synapses onto CCAP neurons (Supplementary Table 3). This connectivity suggests that CCAP neurons are part of the broad network that receives ISN input (Fig 5H). Moreover, the reciprocal regulation of sugar and water ingestion by CCAP neurons argues that multiple peptidergic neurons downstream of the ISNs cooperate to coordinate ingestion of sugar versus water based on specific need.

## DISCUSSION

In this study, we report that the ISNs communicate hunger and thirst states to a complex neural network that reaches several brain regions to regulate sugar and water ingestion (Fig 6). The ISNs synapse with neurons that project to higher brain neuroendocrine centers, including BiT, a novel neuron that reciprocally regulates sugar and water ingestion. Several peptide-releasing and peptide-sensing neurons known to regulate feeding behavior also receive ISN signals, providing the capacity to integrate hunger and thirst signals with many internal signals of nutritional need. These peptidergic neurons, connected to the ISNs via interneurons, contribute differentially to ingestion of sugar and water, with IPC and CCAP neurons reciprocally regulating sugar and water ingestion and CCHa2R-RA neurons modulating water ingestion. Thus, our work argues that the coordinated regulation of a peptidergic network weighs nutrient needs to generate nutrient-specific ingestion.

**Figure 6:**
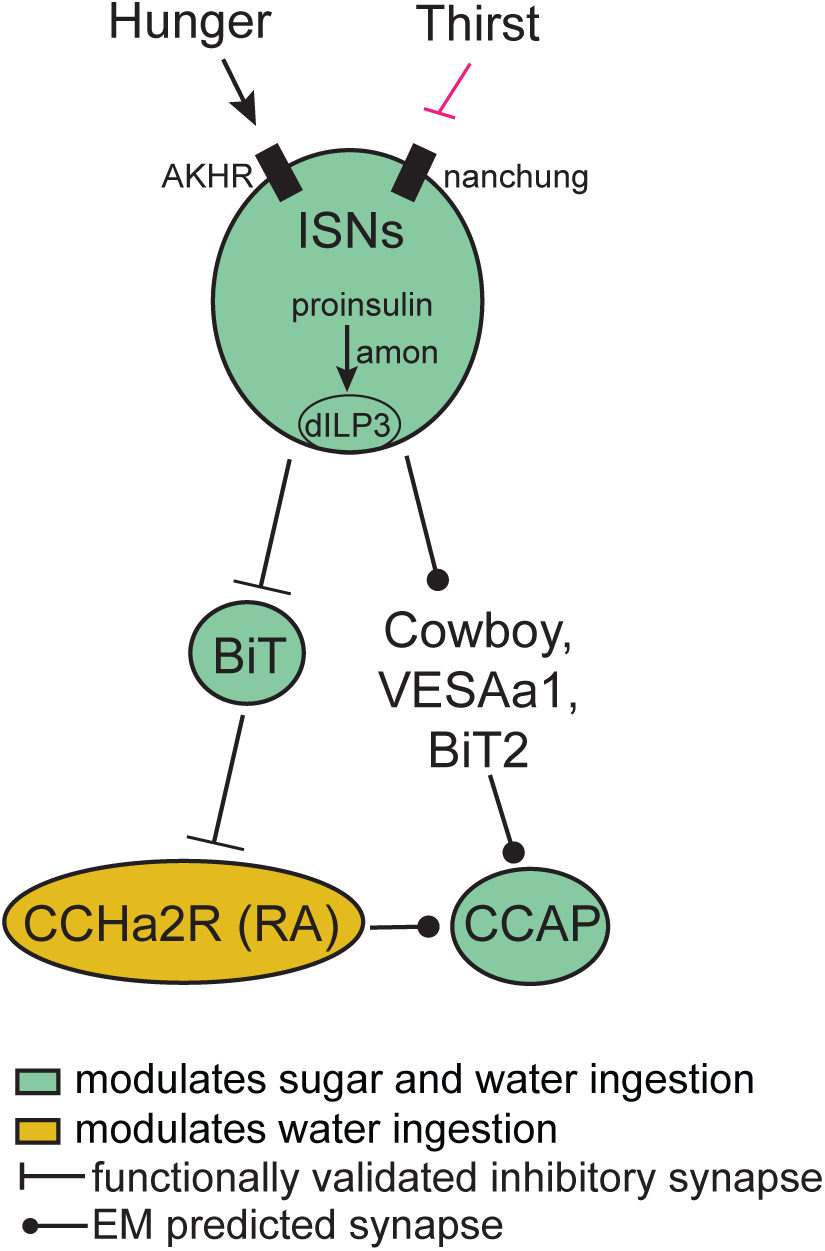
ISN regulation of sugar and water ingestion model. Hunger signals activate the ISN while thirst signals inhibit the ISNs. ISNs use dILP3 as a neurotransmitter and require amontillado (amon) for neuropeptide processing. ISN activity inhibits BiT, which in turn inhibits CCHa2R-RA neurons. CCAP neurons are downstream of the ISNs, connected via Cowboy, VESAa1, BiT2 and CCHa2R-RA neurons. BiT activity inhibits sugar ingestion and promotes water ingestion. CCAP activity promotes sugar modulates sugar and water ingestion modulates water ingestion functionally validated inhibitory synapse EM predicted synapse ingestion and inhibits water ingestion. CCHa2R-RA activity inhibits water ingestion.

### The ISNs influence activity of several brain regions involved in feeding and nutrient homeostasis to coordinate sugar and water ingestion

Previous studies showed that the ISNs sense the hunger signal AKH and changes in hemolymph osmolality associated with thirst to correspondingly alter ISN neural activity. Increased ISN activity promotes sugar ingestion and decreases water ingestion, and decreased ISN activity decreases sugar ingestion and increases water ingestion (Jourjine, Mullaney et al., 2016). Here, we investigated how ISN activity reciprocally regulates sugar and water ingestion according to internal needs by examining the neural network modulated by the ISNs.

We found that the ISNs are predicted to synapse with 100 neurons, including projection neurons that arborize in neuroendocrine centers, SEZ interneurons, and ascending and descending neurons that likely innervate the ventral nerve cord. The majority of the ISN predicted synaptic partners are projection neurons that send arbors via the median bundle to the SMP, a neuroendocrine center (Hartenstein, 2006). This includes the cell type BiT characterized in this study that reciprocally regulates sugar and water ingestion. Local SEZ neurons downstream of the ISNs include DSOG1, which are GABAergic and inhibit consumption (Pool et al., 2014), consistent with the notion that ISN activity directly influences feeding motor programs. In addition, eight uncharacterized descending neurons are downstream of the ISNs, suggesting that they may coordinate feeding with other motor behaviors, such as locomotion or digestion. While the number of ISN postsynaptic partners precludes comprehensive functional and behavioral analysis, the restricted number of brain regions that are direct targets of the ISNs (SMP, SEZ, and possibly ventral nerve cord) is consistent with ISN activity directly regulating neuroendocrine centers and feeding behavior.

We characterized the pathway from the second-order BiT projection neuron that oppositely regulates sugar and water consumption. We found that BiT has 93 predicted synaptic partners, including IPCs which are known to modulate food intake (Nässel & Zandawala, 2020), FLAa3/Lgr3 which have been implicated in regulating ingestion (Laturney et al., 2022, Meissner et al., 2016, Nässel et al, 2013, Yeom et al., 2021), and neurons labeled by the CCHa2R-RA-Gal4 (Deng et al., 2019) which we found regulate water ingestion. BiT downstream neurons innervate several neuropils including the SEZ, SMP, SLP, fan shaped body, and lobula. Therefore, hunger and thirst signals sensed by the ISNs fan out to modulate multiple brain regions via BiT. We speculate that the broad reach of the ISNs serves to modulate different behaviors such as sleep, reproduction, and locomotion based on the hunger or thirst state of the fly.

### Communication between peptidergic neurons coordinates ingestion

Our studies demonstrate that multiple peptidergic neurons participate in regulation of sugar and water ingestion. We find that dILP3 RNAi or *amontillado* RNAi expression in the ISNs recapitulates the ISN loss-of function phenotype, arguing that the ISNs themselves are peptidergic and utilize dILP3 as the neurotransmitter that conveys hunger and thirst signals. The ISNs have increased activity upon AKH detection or low osmolality (hunger signals) (Jourjine, Mullaney et al., 2016), arguing that increased dILP3 release from the ISNs drives sucrose ingestion and limits water ingestion in hungry flies to maintain homeostasis. This conversion of an AKH signal to a dILP3 signal resembles findings in *Drosophila* larvae, where circulating AKH binds to the AKH receptor on IPCs to release dILP3 and promote sucrose consumption (Kim & Neufeld, 2015, Palovcik et al., 1984).

The ISNs modulate activity in many neuroendocrine cells, potentially causing widespread changes in peptide release (Nässel & Zandawala, 2022, Schlegel et al., 2016). We find that ISN activation increases activity of CCAP neurons and CCHa2R-RA neurons, and BiT activation decreases the activity of IPCs. CCAP neurons are orexigenic and communicate to CCAP receptor cells, including IPCs (Zhang et al., 2022) and a subpopulation of neuropeptide F (NPF) neurons (Williams et al., 2020). While this is the first study that characterizes the CCHa2R-RA neurons, the knockin Gal4 line that labels the CCHa2R-RA neurons was generated for the RA isoform of CCHa2 receptor, suggesting that these neurons respond to CCHa2, a peptide produced in the midgut and brain that increases appetite (Deng et al., 2019, Ida et al., 2012, Reiher et al., 2011, Ren et al., 2015). Therefore, CCHa2R-RA neurons potentially integrate the hunger and thirst signals from the ISNs with CCHa2 signals from the gut. IPCs are central regulators of appetite and metabolism, receive multiple direct and indirect signals of nutrient status, and release dILP2, dILP3, and dILP5 (Nässel & Zandawala, 2020). Our finding that the ISNs communicate with multiple peptidergic systems argues that hunger and thirst signals sensed by the ISNs are integrated with other nutritive state signals sensed by ISN downstream neurons for a global assessment of the current nutritional demands of the animal.

### Sugar and water ingestion remain coordinated downstream of the ISNs

Multiple neurons downstream of the ISNs bidirectionally regulate both sugar and water ingestion, arguing that they bias ingestion based on nutrient need. By studying the activation and silencing phenotypes associated with CCAP neurons, we show that acute activation promotes sugar ingestion and limits water ingestion, while silencing these neurons has the opposite effects. These findings are consistent with and expand upon previous studies showing that CCAP neurons promote feeding (Selcho et al, 2018, Williams et al., 2020). IPCs have a more complex role in regulating ingestion, with several studies showing that their acute activation limits ingestion of sucrose or food (Nässel et al., 2015, Semaniuk and Gospodaryov et al., 2018, Wang et al., 2020) and other studies suggesting the opposite (Sudhakar et al., 2020). We find that under the specific conditions of our assay, acute activation of IPCs promotes sucrose ingestion and limits water ingestion. We suspect that differing findings upon IPC manipulation may stem from differences in the deprivation state of the fly, the behavioral assay, the type and timing of neural manipulation, and the food source. As IPCs receive multiple internal state signals, it is possible that activation phenotypes depend on the current state of IPC modulation set by the internal state of the fly.

Overall, our work sheds light on neural circuit mechanisms that translate internal nutrient abundance cues into the coordinated regulation of sugar and water ingestion. We show that the hunger and thirst signals detected by the ISNs influence a network of peptidergic neurons that act in concert to prioritize ingestion of specific nutrients based on internal needs. We hypothesize that multiple internal state signals are integrated in higher brain regions such that combinations of peptides and their actions signify specific needs to drive ingestion of appropriate nutrients. As peptide signals may act at a distance and may cause long-lasting neural activity state changes, studying their integration over space and time is a future challenge to further illuminate homeostatic feeding regulation.

## MATERIALS AND METHODS

| Key resources table |  |  |
| --- | --- | --- |
| Drosophila strains | Source or Reference | Identifier |
| UAS-nSynaptobrevin RNAi | Bloomington Drosophila Stock Center | BDSC 31983 |
| UAS-dcr2 | Bloomington Drosophila Stock Center | BDSC 24650 |
| UAS-Trh RNAi | Bloomington Drosophila Stock Center | BDSC 25842 |
| UAS-ChAT RNAi | Bloomington Drosophila Stock Center | BDSC 25856 |
| UAS-Tbh RNAi | Bloomington Drosophila Stock Center | BDSC 27667 |
| UAS-Hdc RNAi | Bloomington Drosophila Stock Center | BDSC 26000 |
| UAS-VMAT RNAi | Bloomington Drosophila Stock Center | BDSC 31257 |
| UAS-GAD1 RNAi | Bloomington Drosophila Stock Center | BDSC 28079 |
| UAS-DDC RNAi | Bloomington Drosophila Stock Center | BDSC 27030 |
| UAS-DVGlut RNAi | Bloomington Drosophila Stock Center | BDSC 27538 |
| UAS-sNPF RNAi | Bloomington Drosophila Stock Center | BDSC 25867 |
| UAS-VGAT RNAi | Bloomington Drosophila Stock Center | BDSC 41958 |
| UAS-TDC2 RNAi | Bloomington Drosophila Stock Center | BDSC 25871 |
| UAS-dILP1 RNAi | Bloomington Drosophila Stock Center | BDSC 32861 |
| UAS-dILP2 RNAi | Bloomington Drosophila Stock Center | BSC 32475 |
| UAS-dILP3 RNAi | Bloomington Drosophila Stock Center | BSC 31492 |
| UAS-dILP4 RNAi | Bloomington Drosophila Stock Center | BDSC 33682 |
| UAS-dILP5 RNAi | Bloomington Drosophila Stock Center | BDSC 31378 |
| UAS-dILP6 RNAi | Bloomington Drosophila Stock Center | BDSC 33684 |
| UAS-dILP7 RNAi | Bloomington Drosophila Stock Center | BDSC 32862 |
| UAS-amon RNAi | Bloomington Drosophila Stock Center | BDSC 29009 |
| ISN-Gal4 (VT011155-Gal4) | FlyLight, Janelia Research Campus | Fly Light ID 54404 |
| ISN-LexA (GMR34G02-LexA) | Bloomington Drosophila Stock Center | BDSC 54138 |
| UAS-myrGFP.QUAS-mtdTomato-3xHA; trans-Tango | Bloomington Drosophila Stock Center | BDSC 77124 |
| VT002073-Gal4.AD | Bloomington Drosophila Stock Center | BDSC 71871 |
| VT040568-Gal4.DBD | Bloomington Drosophila Stock Center | BDSC 72902 |
| UAS-csChrimson.mVenus | Bloomington Drosophila Stock Center | BDSC 55134 |
| LexAop-ChrimsonR.mCherry | Gift from Jayaraman Lab |  |
| UAS-ArcLight | Bloomington Drosophila Stock Center | BDSC 51056 |
| Empty split | Bloomington Drosophila Stock Center | BDSC 79603 |
| ppk28-Gal4 | Cameron et al 2010. | BDSC 93020 |
| Gr5a-Gal4 | Chyb et al 2003. | BDSC 57592, 57591 |
| CCha2R-RA-Gal4 | Bloomington Drosophila Stock Center | BDSC 84603 |
| LexAop-CsChrimson.tdTomato (III) | Bloomington Drosophila Stock Center | BDSC 82183 |
| UAS-GCaMP6s (III) | Bloomington Drosophila Stock Center | BDSC 42749 |
| 20XUAS-GCaMP7b | Bloomington Drosophila Stock Center | BDSC 79029 |
| CCAP-Gal4 (II) | Bloomington Drosophila Stock Center | BDSC 25685 |
| CCAP-Gal4 (III) | Bloomington Drosophila Stock Center | BDSC 25686 |
| CCha2R-RA-LexA | Bloomington Drosophila Stock Center | BDSC 84363 |
| dILP2-LexA | Li and Gong 2015. |  |

### Fly husbandry

All experiments and screening were carried out with adult *D. melanogaster* females reared on standard cornmeal-agar-molasses medium, at 25°C, 65-70% humidity, on a 12 hr light: 12 hr dark cycle. Flies used in optogenetic assays were reared on food containing 0.25mM all-trans-retinal (Sigma-AldrichSigna-Aldrich) in darkness, before and after eclosion.

### Temporal consumption assay (TCA)

Flies were anesthetized using CO_2_ and then fixed to a glass slide with nail polish. Flies recovered for 2 hours in a humidified box if testing for sucrose ingestion, or in a desiccated box with Drierite if testing for water ingestion. Immediately before testing for sucrose ingestion, flies were given water until they no longer responded to 3 consecutive water presentations. In testing, flies were presented with the tastant (water or 1M sucrose) 10 times and consumption time was manually recorded. All experiments were done in a dark, temperature- and humidity-controlled room. IR lights and IR cameras were used to conduct experiments in the dark. All water tests were done in the morning and all sugar tests were done in the afternoon. For optogenetic activation experiments, we expressed the light activated ion channel Chrimson in the neurons of interest and activated these neurons using a 635nm laser (Laserglow). For silencing experiments, we expressed RNAi against nSynaptobrevin in neurons of interest.

### *In vivo* calcium imaging

Calcium imaging studies were carried out as described in Shiu, Sterne et al. (2022). Mated female flies were dissected for calcium imaging studies 5-14 days post-eclosion. Flies were briefly anesthetized with ice and placed in a custom plastic holder at the neck to isolate the head from the rest of the body. The head was then immobilized using UV glue, the proboscis was immobilized using wax, and the esophagus was cut to provide unobstructed imaging access to the SEZ. All flies imaged were sated. *In vivo* calcium imaging with optogenetic activation was performed in a 2-photon microscope using a Scientifica Hyperscope with resonant scanning, a piezo drive, and a 20× water immersion objective (NA = 1.0) with 1.8-3× digital zoom, depending on the cell type imaged. Calcium responses were recorded with a 920 nm laser and optogenetic stimulation was achieved with a 660 nm LED. 2s LED stimulation paradigm: 20s off, 2s on, 30s off, 2s on, 30s off. 30s LED stimulation paradigm: 20s off, (1s on, 1s off) × 15, 60s off. For the 2s LED stimulation, 80 stacks of 20 z slices of 4-5 μm were acquired at 0.667 Hz. For the 30s stimulation, 125 stacks of 20 z slices of 4-5 μm were acquired at 0.667 Hz. Analysis was done on max-z projections of the 20 z slices. %ΔF/F = 100*((Ft-F0)/F0), where Ft is the fluorescence of the Neuron ROI - the Background ROI at each timepoint and F0 is the mean Ft for the 23 time points prior to stimulus onset. Quantification was carried out in GraphPad Prism. A mean fluorescence intensity for LED off and LED on was calculated for each fly. For the 2s LED stimulation, mean intensity for LED off was calculated for 5 timepoints immediately before LED exposure and mean intensity for LED on was calculated for 5 timepoints during LED exposure. For the 30s stimulation, mean intensity for LED off was calculated for 28 timepoints immediately before LED exposure and mean intensity for LED on was calculated for 28 timepoints during LED exposure. Paired t-test or paired Wilcoxon test was performed. ROI for CCHa2R-RA imaging was CCHa2R-RA neurites in SEZ. ROI for CCAP imaging was CCAP neurites. ROI for IPC imaging was all IPC somas.

### *In vivo* voltage imaging

Voltage imaging studies were carried out exactly as calcium imaging studies described above. ROI for BiT imaging was BiT soma.

### Immunohistochemistry

All brain and CNS dissections and immunostaining (unless directly addressed) were carried out as described (https://www.janelia.org/project-team/flylight/protocols, ‘IHC-Anti-GFP’) substituting the below antibodies and eschewing the pre-embedding fixation steps. Ethanol dehydration and DPX mounting was carried out as described (https://www.janelia.org/project-team/flylight/protocols, ‘DPX Mounting’).

Primary antibodies:

- mouse α-Brp (nc82, DSHB, University of Iowa, USA) at 1:40
- chicken α-GFP (Invitrogen, A10262) at 1:1000
- rabbit α-dsRed (Takara, Living Colors 632496) at 1:1000

Secondary antibodies:

- goat α-mouse AF647 (Invitrogen, A21236) at 1:500
- goat α-chicken AF488 (Life Technologies, A11039) at 1:1000
- goat α-rabbit AF568 (Invitrogen, A21236) at 1:1000

Images were acquired with a Zeiss LSM 880 NLO AxioExaminer with Airyscan and Coherent Chameleon Vision or Zeiss LSM 780 Laser Scanning Confocal Microscope at the Berkeley Molecular Imaging Center with a Plan-Apochromat 20×/1.0 W, 40× W, 40×/1.4 oil, or 63×/1.4 oil objective. Images were prepared in Fiji.

### Electron microscopy neural reconstructions and connectivity

Neurons were reconstructed in a serial section transmission electron volume (Full Adult Female Brain, Zheng and Lauritzen et al., 2018) using the CATMAID software (Saalfeld et al., 2009). Fully manual reconstructions were generated by following the branches of the neuron and marking the center of each branch, thereby creating a ‘skeleton’ of each neuron. In addition to fully manual reconstructions, segments of an automated segmentation (Li et al., 2019) were proofread and expanded to generate complete reconstructions. In addition to the skeleton tracing, new chemical synapses were also annotated as previously described (Zheng and Lauritzen et al. 2018). Downstream synaptic targets of the ISNs and BiT were then traced out from these additional locations using both manual and assisted tracing techniques as described above. Neurons traced in CATMAID, including ISNs and BiT, were all located in Flywire (flywire.ai), which uses the same EM electron microscopy dataset (Zheng and Lauritzen et al. 2018). To identify synaptic partners, we used connectome annotation versioning engine (CAVE, Buhmann et al. 2021, Heinrich et al. 2018) using a cleft score cutoff of 50 to generate synapses of relatively high confidence (Heinrich et al. 2018; Baker et al., 2022). FAFB neural reconstructions were visualized using NAVis (Copyright 2018, Philipp Schlegel), which is based on natverse (Bates et al., 2020).

### BiT split-Gal4 generation

We created a color depth max intensity projection (CDM) mask of BiT reconstructed EM skeleton and used CDM mask searching (Otsuna et al., 2018) to find enhancers whose expression patterns seemed to include the desired cell type using MCFO (Nern et al., 2015) screening of subsets of the Janelia Research Campus and Vienna Tile Gal4 collections. Construction of stable split-Gal4 lines was performed as previously described (Dionne et al., 2018, Sterne et al., 2020). Immunohistochemistry and confocal imaging was used to determine successful split-Gal4 combinations.

### Identification of GAL4 lines from EM reconstructions

Visual inspection of Gal4 collections was used to determine cell type. Images of potential Gal4 lines were skeletonized in FIJI, converted into .swc format using natverse (Bates et al., 2021), and uploaded to Flywire using the Flywire Gateway. This generated pointclouds that were used to identify the neurons of interest. As Flywire permits exhaustive searching of neurons in an area, we examined all neurons in the region of interest to conclusively identify our neuron of interest.

### Statistical analysis

Statistical tests were performed in GraphPad Prism. For all group comparisons, data was first tested for normality with the KS normality test (alpha = 0.05). If all groups passed then groups were compared with a parametric test, but if at least one group did not pass, groups were compared with a non-parametric version. All statistical tests, significance levels, and number of data points (N) are specified in the figure legend. All datasets from optogenetic behavior assays were normalized within each genotype. To generate this normalized dataset, data from females within the no light condition was averaged, creating a “no-light mean” for each genotype. This value was subtracted from each individual female within the light condition of the corresponding genotype. This dataset was then graphed, and statistical analyses were performed as outlined above.

## ACKNOWLEDGEMENTS

Members of the Scott lab provided contributions to experimental design, data analysis, and manuscript preparation. This work was supported by NIH R01GM128209 (K.S.), R01GM128209 Diversity Supplement (A.G.S.), UC LEADs fellowship (A.D.T.) and International Fellowship for CONICET Researchers (G.P.). Neuronal reconstruction for this project took place in a collaborative CATMAID environment in which 27 labs are participating to build connectomes for specific circuits. Development and administration of the FAFB tracing environment and analysis tools were funded in part by National Institutes of Health BRAIN Initiative grant 1RF1MH120679-01 to Davi Bock and Greg Jefferis, with software development effort and administrative support provided by Tom Kazimiers (Kazmos GmbH) and Eric Perlman (Yikes LLC). We thank Peter Li, Viren Jain and colleagues at Google Research for sharing the automatic segmentation (Li et al., 2019). Tracing in Cambridge was supported by Wellcome Trust (203261/Z/16/Z) and ERC (649111) awards to G. Jefferis. Neurons were also reconstructed and proofread in FlyWire, where we also identified pre- and postsynaptic partners. We acknowledge the Princeton FlyWire team and members of the Murthy and Seung labs for development and maintenance of FlyWire (supported by BRAIN initiative grant MH117815 to Murthy and Seung). In addition to our tracing efforts in CATMAID and proofreading in FlyWire, the following labs greatly contributed to the proofreading in FlyWire of the neurons characterized in this study: Murthy and Seung Labs (76.06%), Jefferis Lab (13.57%) and Jinseop Kim Lab (7.21%). We are appreciative of the proofreading contributed by other labs including the Anderson, Bock, Dacks, Dickson, Huetteroth, Pankratz, Seeds/Hampel, Selcho, Simpson, Waddell, Wilson and Wolf Labs, and Janelia tracers (Supplementary Table 1, Supplementary Table 2, Supplementary Table 3). Vivek Jayaraman provided unpublished fly lines used in this study. We thank Stefanie Engert for tracing two ISNs in CATMAID, Zepeng Yao for identifying CCHa2R-RA neurons, Phil Shiu for identifying DSOG1 neurons, and Amanda Abusaif for her tracing and proofreading efforts. Confocal imaging experiments were conducted at the CRL Molecular Imaging Center, RRID:SCR_017852, supported by NSF DBI-1041078 and the Helen Wills Neuroscience Institute. We thank Holly Aaron and Feather Ives for microscopy training and support.

## AUTHOR CONTRIBUTIONS

A.G.S. and K.S. conceived and designed the study and wrote the manuscript. A.G.S performed the FAFB circuit building and cell identification, BiT split-Gal4 creation, functional imaging experiments, immunohistochemistry, and data analysis. G.P. performed behavior experiments and BiT split-Gal4 creation. N.J performed the ISN neurotransmitter RNAi screen. A.D.T. traced ISNs and BiT in CATMAID.

## COMPETING INTERESTS

The authors declare no competing interests.

## TABLES

**Table S1:**
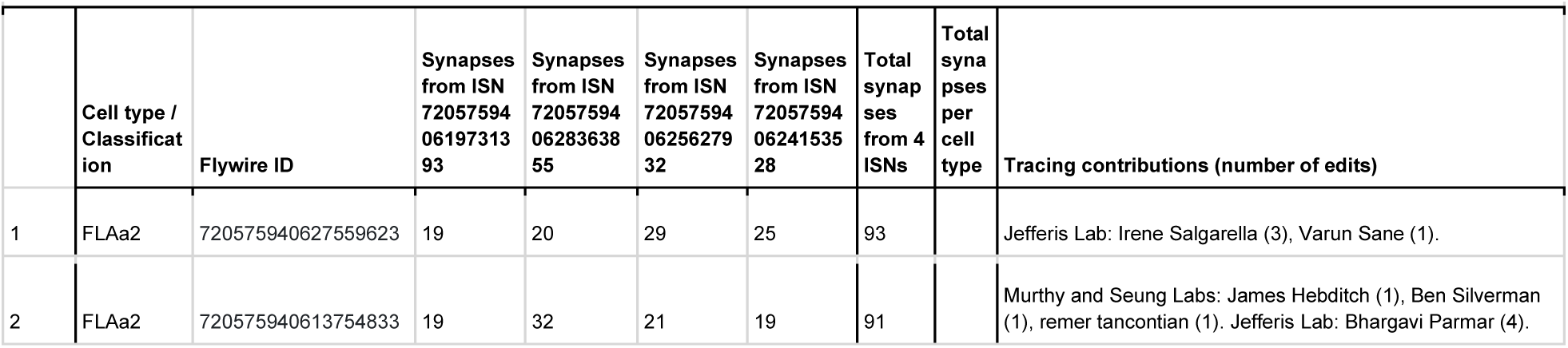
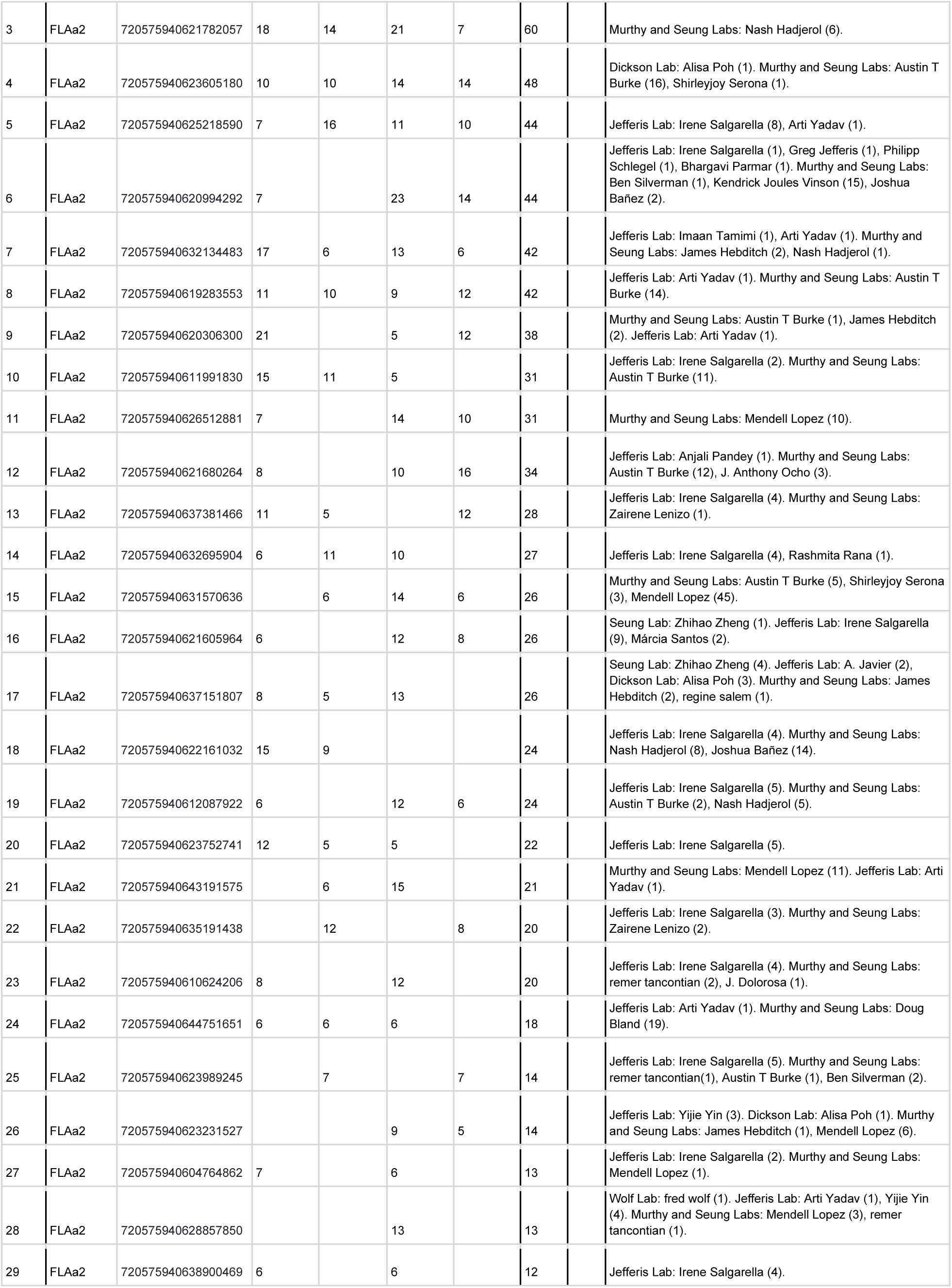
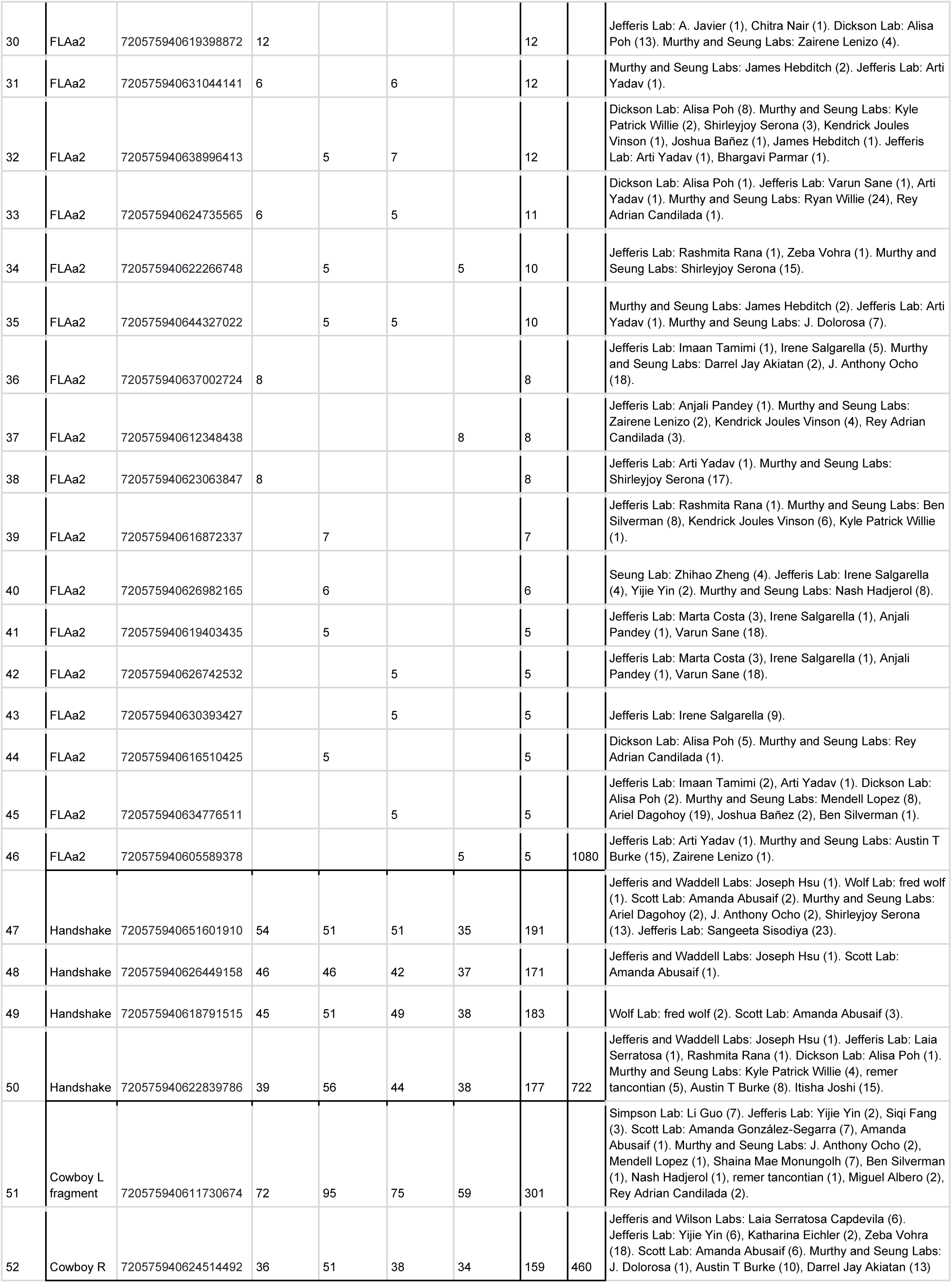
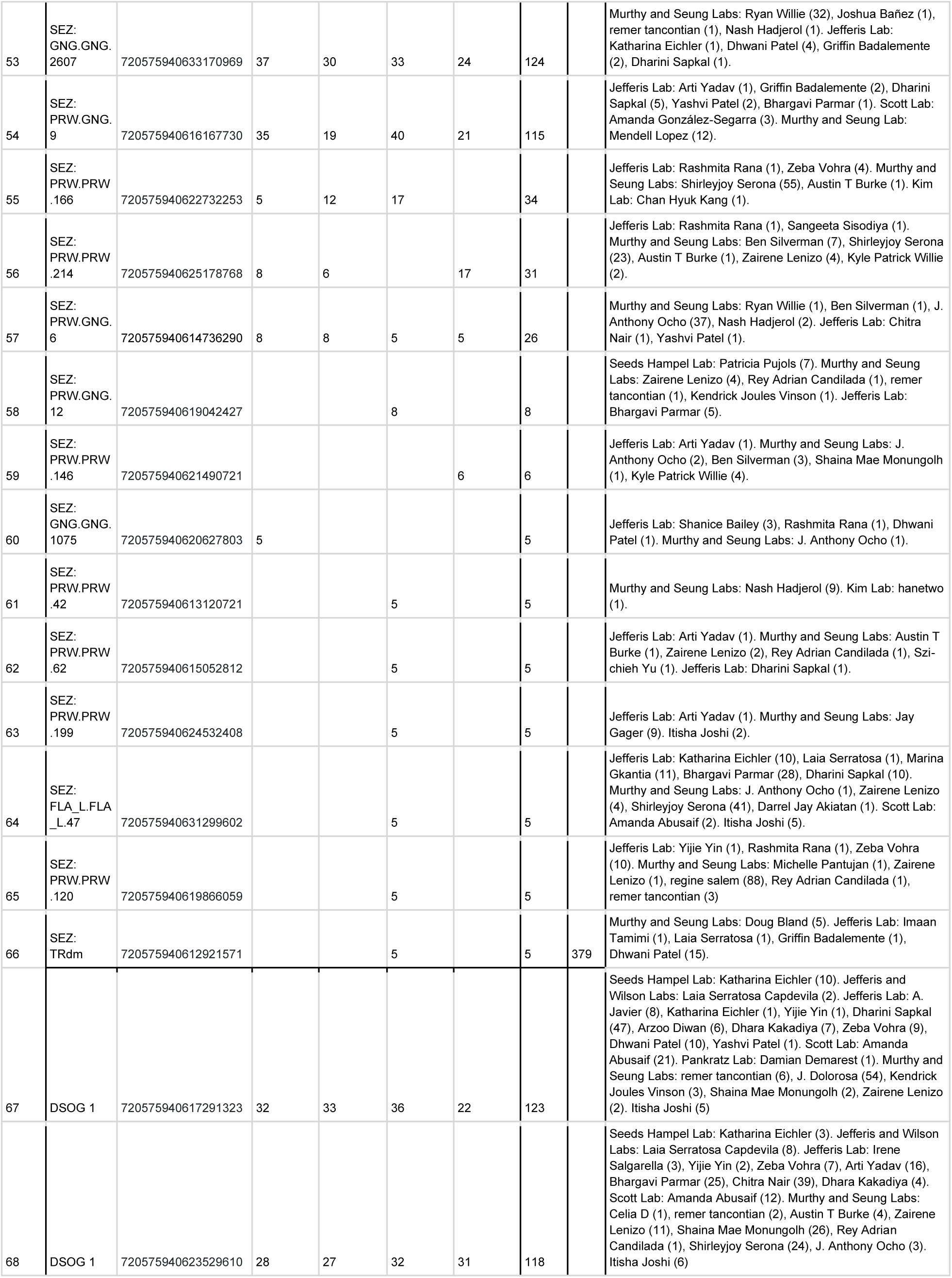
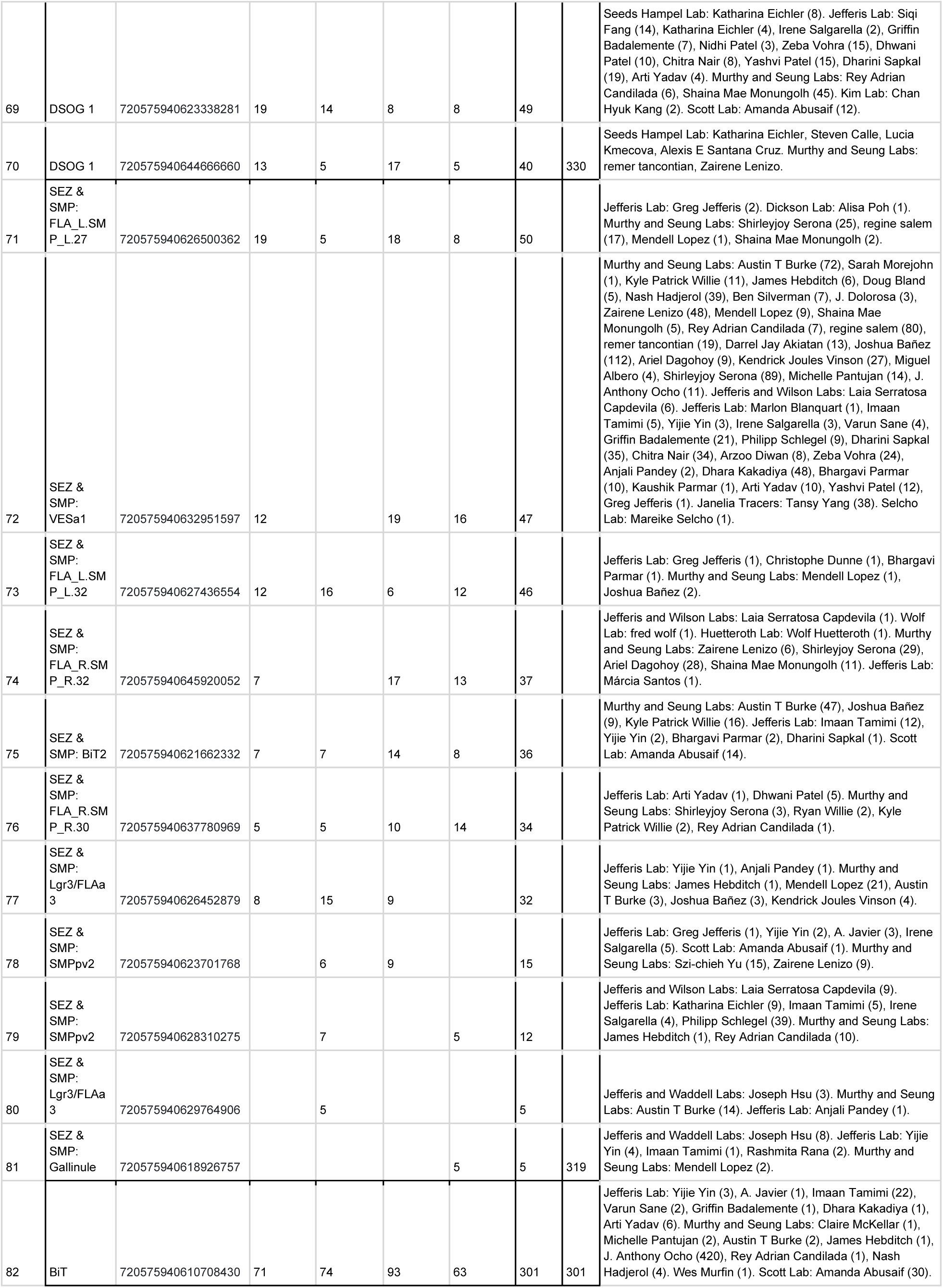
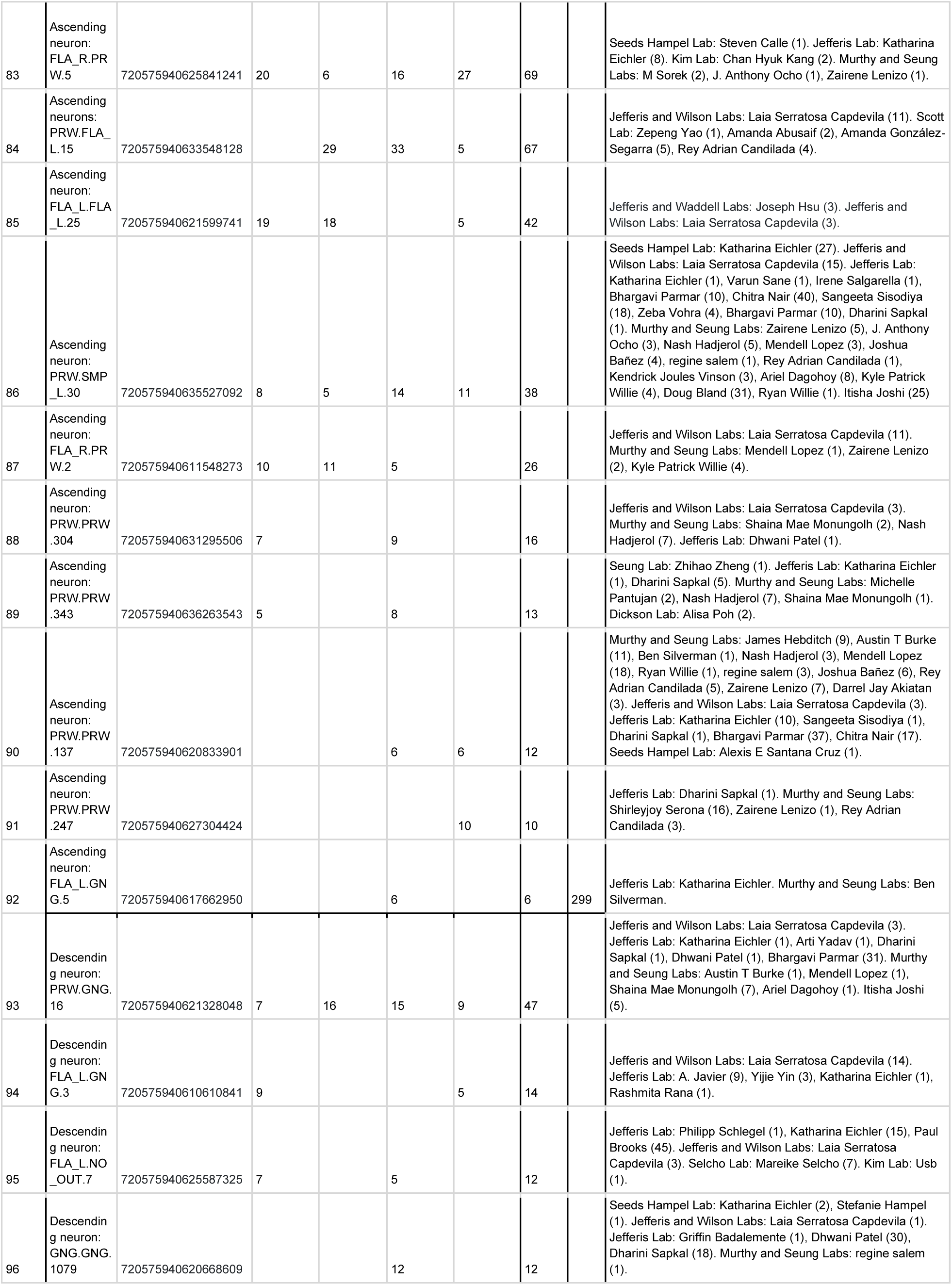
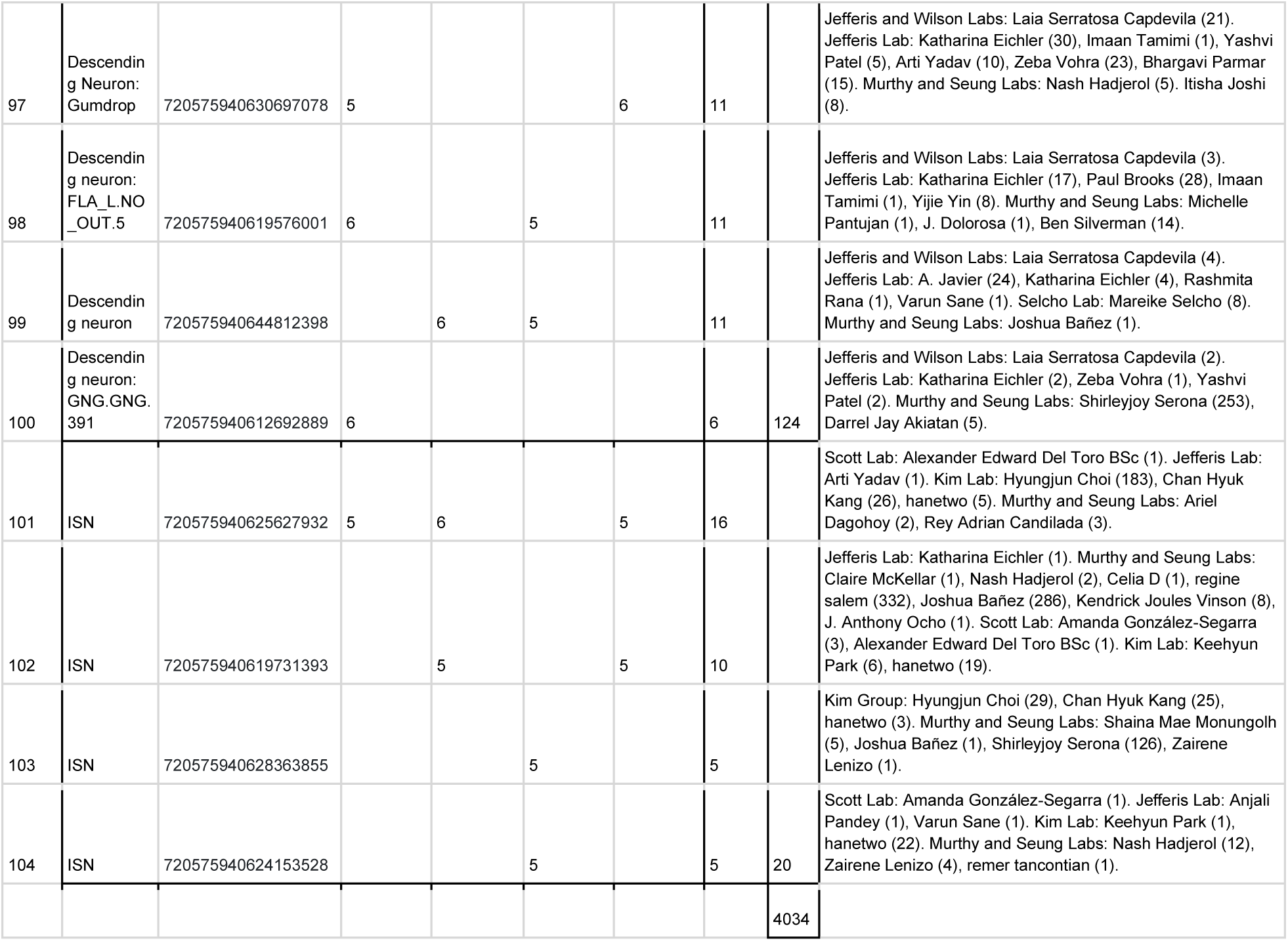
ISN postsynaptic partners.

**Table S2:** BiT postsynaptic neurons.

| Table S2: BiT postsynaptic neurons |  |  |  |  |  |
| --- | --- | --- | --- | --- | --- |
|  | Name | Neuron ID | Synapses from BiT | Total synapses from BiT per cell type | Tracing contributions (number of edits) |
| 1 | IPC | 720575940650527222 | 54 |  | Jefferis Lab: Imaan Tamimi (2), Anjali Pandey (1). Murthy and Seung Labs: Austin T Burke (4). |
| 2 | IPC | 720575940603765280 | 44 |  | Maimon Lab: Gaby Maimon (1). Jefferis Lab: Katharina Eichler (2). Murthy and Seung Labs: Austin T Burke (3). |
| 3 | IPC | 720575940620932045 | 39 |  | Jefferis Lab: Imaan Tamimi (2), Laia Serratos (7). Murthy and Seung Labs: Austin T Burke (6). |
| 4 | IPC | 720575940612923390 | 33 |  | Jefferis Lab: Imaan Tamimi (1), Laia Serratos (6), Markus Pleijzier (1), Rashmita Rana (1). Murthy and Seung Labs: Ryan Willie (1). |
| 5 | IPC | 720575940625379859 | 30 |  | Jefferis Lab: A. Javier (4), Irene Salgarella (2), Laia Serratos (9). Murthy and Seung Labs: Austin T Burke (4). |
| 6 | IPC | 720575940622897639 | 28 |  | Seung Lab: Zhihao Zheng (1), Jefferis Lab: A. Javier (1). Dacks Lab: Andrew Dacks (1). Murthy and Seung Labs: Austin T Burke (7). |
| 7 | IPC | 720575940643539566 | 26 |  | Jefferis Lab: Imaan Tamimi (3), A. Javier (7), Laia Serratosa (4), Rashmita Rana (1). Murthy and Seung Labs: Austin T Burke (13). |
| 8 | IPC | 720575940620628957 | 25 |  | Jefferis Lab: Imaan Tamimi (1), A. Javier (6). Murthy and Seung Labs: Austin T Burke. (18) |
| 9 | IPC | 720575940623081400 | 22 |  | Jefferis Lab: Markus Pleijzier (5). Murthy and Seung Labs: Austin T Burke (18). |
| 10 | IPC | 720575940631884883 | 22 |  | Dacks Lab: Andrew Dacks (1). Jefferis Lab: Imaan Tamimi (1), Laia Serratosa (8). Murthy and Seung Labs: Austin T Burke (7). Itisha Joshi (2). |
| 11 | IPC | 720575940628363820 | 22 |  | Jefferis Lab: Imaan Tamimi (1). Dacks Lab: Andrew Dacks (1). Murthy and Seung Labs: Austin T Burke (6). Scott Lab: Meghan Laturney (1). |
| 12 | IPC | 720575940624064295 | 22 |  | Dacks Lab: Andrew Dacks (1). Jefferis Lab: Imaan Tamimi (3), Laia Serratosa (10), Anjali Pandey (1). Murthy and Seung Labs: Austin T Burke (5). |
| 13 | IPC | 720575940628199802 | 21 |  | Jefferis Lab: Markus Pleijzier (1), A. Javier (1), Laia Serratosa (9), Philipp Schlegel (1). Murthy and Seung Labs: Austin T Burke (1). Wolf Lab: Fred Wolf (1). |
| 14 | IPC | 720575940623586284 | 17 |  | Jefferis Lab: Imaan Tamimi (1), Márcia Santos (4). Murthy and Seung Labs: Austin T Burke (66). |
| 15 | IPC | 720575940618694827 | 14 |  | Jefferis Lab: A. Javier (3), Imaan Tamimi (1), Laia Serratosa (12), Anjali Pandey (1). |
| 16 | IPC | 720575940614623455 | 10 |  | Murthy and Seung Labs: James Hebditch (4), Nash Hadjerol (1), Doug Bland (6), Joshua Bañez (6). Jefferis Lab: Philipp Schlegel (1). |
| 17 | IPC | 720575940611254681 | 7 |  | Jefferis Lab: Markus Pleijzier (2), A. Javier (3), Imaan Tamimi (9). Murthy and Seung Labs: Austin T Burke (13). Kim Lab: Minsik Yun (2). |
| 18 | IPC | 720575940615911380 | 6 | 442 | Jefferis Lab: A. Javier (6), Laia Serratosa (6), Philipp Schlegel (1). Murthy and Seung Labs: Ryan Willie (1). |
| 19 | Lgr3/FLAa3 | 720575940626983185 | 36 |  | Jefferis Lab: Irene Salgarella (10). Murthy and Seung Labs: James Hebditch (5), Kyle Patrick Willie (4), Rey Adrian Candilada (1), Kendrick Joules Vinson (11), Zairene Lenizo (2), Nash Hadjerol (4). |
| 20 | Lgr3/FLAa3 | 720575940620201084 | 32 |  | Jefferis Lab: Irene Salgarella (7), Laia Serratosa (4). Murthy and Seung Labs: James Hebditch (4). |
| 21 | Lgr3/FLAa3 | 720575940613895934 | 31 |  | Jefferis Lab: Irene Salgarella (10), Dharini Sapkal (2), Dhvani Patel (12). Murthy and Seung Labs: James Hebditch (2), Zairene Lenizo (2), Joshua Bañez (8). |
| 22 | Lgr3/FLAa3 | 720575940629696763 | 30 |  | Jefferis Lab: Anjali Pandey (1). Murthy and Seung Labs: James Hebditch (4), Nash Hadjerol (6), Ben Silverman (1), Darrel Jay Akiatan (2). |
| 23 | Lgr3/FLAa3 | 720575940644053271 | 19 |  | Jefferis Lab: Anjali Pandey (1), Christopher Dunne (11). Murthy and Seung Labs: Zairene Lenizo (1). |
| 24 | Lgr3/FLAa3 | 720575940630329903 | 19 |  | Jefferis Lab: Irene Salgarella (15). Murthy and Seung Labs: James Hebditch (3). |
| 25 | Lgr3/FLAa3 | 720575940616987426 | 19 |  | Murthy and Seung Labs: James Hebditch (4). Jefferis Lab: Anjali Pandey (8), Christopher Dunne (4), Sangeeta Sisodiya (1), Dharini Sapkal (4). |
| 26 | Lgr3/FLAa3 | 720575940639420032 | 19 |  | Jefferis Lab: Irene Salgarella (16). Murthy and Seung Labs: James Hebditch (4), Zairene Lenizo (1). |
| 27 | Lgr3/FLAa3 | 720575940614714299 | 18 |  | Jefferis Lab: Irene Salgarella (7). Murthy and Seung Labs: Nash Hadjerol (1). |
| 28 | Lgr3/FLAa3 | 720575940632879842 | 18 |  | Scott Lab: Zepeng Yao (17). Jefferis Lab: Arti Yadav (1), Bhargavi Parmar (6). Murthy and Seung Labs: J. Anthony Ocho (1), Austin T Burke (1), Zairene Lenizo (2). |
| 29 | Lgr3/FLAa3 | 720575940626179658 | 15 |  | Jefferis Lab: Irene Salgarella (10). |
| 30 | Lgr3/FLAa3 | 720575940626327070 | 15 | 271 | Murthy and Seung Labs: Austin T Burke (1), James Hebditch (4), Ariel Dagohoy (4), Nash Hadjerol (2). Janelia tracers: Tansy Yang (2). Jefferis Lab: Irene Salgarella (8), Laia Serratosa (3). Scott Lab: Zepeng Yao (3). Pankratz Lab: Damian Demarest (2). |
| 31 | SMP & SLP:<br>SMP_L.SMP_L.276 | 720575940617650203 | 45 |  | Jefferis and Waddell Labs: Joseph Hsu (2). Murthy and Seung Labs: Kyle Patrick Willie (32). Jefferis Lab: Laia Serratosa (1), Anjali Pandey (1), Yijie Yin (4). |
| 32 | SMP & SLP:<br>SMP_R.SMP_R.27<br>8 | 720575940619352198 | 44 |  | Murthy Lab: Lucas Encarnacion-Rivera (1), Bock Lab: Davi Bock (4). Jefferis Lab: Yijie Yin (13), Greg Jefferis (15). |
| 33 | SMP & SLP:<br>SMPpv1; right | 720575940631373869 | 38 |  | Jefferis and Waddell Labs: Joseph Hsu (2), Jefferis Lab: Irene Salgarella (1). Murthy and Seung Labs: J. Dolorosa (2), Kyle Patrick Willie (4). |
| 34 | SMP & SLP:<br>SMPpv1; left | 720575940633754292 | 26 |  | Jefferis Lab: Yijie Yin (2). Murthy and Seung Labs: Ben Silverman (1). |
| 35 | SMP & SLP:<br>SMP_R.SMP_R.80<br>8 | 720575940635354981 | 12 |  | Murthy and Seung Labs: Zairene Lenizo (1), Nash Hadjerol (24), remer tancontian (5). Jefferis Lab: Anjali Pandey (1). |
| 36 | SMP & SLP:<br>SMPpd1; left | 720575940627327750 | 12 |  | Jefferis Lab: Rashmita Rana (1). Murthy and Seung Labs: James Hebditch (3), remer tancontian (1). |
| 37 | SMP & SLP:<br>SMP_R.SMP_R.70 | 720575940611332722 | 11 |  | Jefferis Lab: Yijie Yin (1), Rashmita Rana (1). Murthy and Seung Labs: Kyle Patrick Willie (10). |
| 38 | SMP & SLP:<br>SMPpd1; left | 720575940619099558 | 10 |  | Jefferis Lab: A. Javier (2), Arti Yadav (1). |
| 39 | SMP & SLP:<br>SMP_R.SMP_R.74<br>2 | 720575940631589407 | 9 |  | Murthy and Seung Labs: Austin T Burke (17), James Hebditch (2), Kendrick Joules Vinson (1). Jefferis Lab: Anjali Pandey (1). |
| 40 | SMP & SLP: SLPal1 | 720575940618140283 | 8 |  | Jefferis Lab: A. Javier (3). Murthy and Seung Labs: Nash Hadjerol (1), Ariel Dagohoy (1), James Hebditch (2). |
| 41 | SMP & SLP:<br>SMPpd2 | 720575940619949556 | 7 |  | Jefferis Lab: Irene Salgarella (3), Anjali Pandey (1). |
| 42 | SMP & SLP:<br>SMP_R.SMP_R.75<br>7 | 720575940632526547 | 6 |  | Murthy and Seung Labs: Austin T Burke (1), Kyle Patrick Willie (6), Mendell Lopez (10), Doug Bland (24), remer tancontian (6). Jefferis Lab: Anjali Pandey (1). |
| 43 | SMP & SLP:<br>SMP_L.SMP_L.706 | 720575940629298679 | 6 |  | Murthy Lab: Lucas Encarnacion-Rivera (3). Jefferis Lab: A. Javier (1), Varun Sane (8), Arti Yadav (1). Murthy and Seung Labs: Kendrick Joules Vinson (7), remer tancontian (2). |
| 44 | SMP & SLP:<br>SMP_R.SMP_R.90<br>6 | 720575940640456923 | 6 |  | Jefferis Lab: Philipp Schlegel (3), Yijie Yin (1), Varun Sane (6), Anjali Pandey (1). Selcho Lab: Mareike Selcho (17). Murthy and Seung Labs: Mendell Lopez (19), Kyle Patrick Willie (1). |
| 45 | SMP & SLP: DM3;<br>right | 720575940621647498 | 5 |  | Jefferis Lab: A. Javier (1), Yijie Yin (1). |
| 46 | SMP & SLP:<br>DM3_canonical | 720575940619899668 | 5 |  | Jefferis Lab: Yijie Yin (16). Murthy and Seung Labs: Nash Hadjerol (1), Kyle Patrick Willie (1). |
| 47 | SMP & SLP:<br>SMP_R.SMP_R.67<br>7 | 720575940629901307 | 5 |  | Jefferis Lab: Yijie Yin (2), Chitra Nair (1), Varun Sane (3). Murthy and Seung Labs: Zairene Lenizo (2). |
| 48 | SMP & SLP:<br>SLP_L.LH_L.5 | 720575940616707545 | 5 |  | Jefferis and Wilson Labs: Laia Serratos Capdevila (1). Wes Murfin (4). Jefferis Lab: Yijie Yin (3), Varun Sane (1), Dhara Kakadiya (2), Chitra Nair (4). Murthy and Seung Labs: Doug Bland (3), Kendrick Joules Vinson (23), Joshua Bañez (10), Rey Adrian Candilada (1). |
| 49 | SMP & SLP:<br>SLP_R.SLP_R.557 | 720575940629163419 | 5 |  | Murthy and Seung Labs: Austin T Burke (3). Jefferis Lab: Tomke S (14), Griffin Badalemente (4), Yijie Yin (1), Varun Sane (4), Dhvani Patel (1). |
| 50 | SMP & SLP:<br>SMP_L.SMP_L.578 | 720575940626063688 | 5 | 270 | Murthy and Seung Labs: Austin T Burke (13). |
| 51 | CCHa2R (RA) | 720575940638190133 | 65 |  | Murthy and Seung Labs: Claire McKellar (1). Jefferis Lab: Anjali Pandey (1), Laia Serratos (117), Dhvani Patel (3). Scott Lab: Amanda Abusaif (1), Zepeng Yao (27). Kim Lab: hanetwo (1). |
| 52 | CCHa2R (RA) | 720575940615181910 | 64 |  | Jefferis Lab: Anjali Pandey (1). Scott Lab: Amanda Abusaif (12). Murthy and Seung Labs: Michelle Pantujan (41), Ryan Willie (459). |
| 53 | CCHa2R (RA) | 720575940629642460 | 50 |  | Jefferis and Waddell Labs: Joseph Hsu (1). Murthy and Seung Labs: Claire McKellar (12), Ryan Willie (1), Shirleyjoy Serona (1), Doug Bland (296), J. Anthony Ocho (3). Jefferis Lab: Arti Yadav (1). Kim Lab: Hyungjun Choi (74). |
| 54 | CCHa2R (RA) | 720575940611411162 | 49 | 228 | Murthy and Seung Labs: Claire McKellar (2), Nash Hadjerol (2), Zairene Lenizo (1). Jefferis Lab: Irene Salgarella (3). Scott Lab: Amanda Abusaif (11). Kim Lab: hanetwo (14), Chan Hyuk Kang (2). |
| 55 | SMP & SEZ:<br>ADM09 | 720575940628462927 | 48 |  | Anderson Lab: Altyn Rymbek (2). Jefferis Lab: Imaan Tamimi (6), Laia Serratos (10), Rashmita Rana (1), Sangeeta Sisodiya (1). Murthy and Seung Labs: James Hebditch (1), Joshua Bañez (1), Mendell Lopez (58), Zairene Lenizo (2), J. Dolorosa (1). Kim Lab: Minsik Yun (1). |
| 56 | SMP & SEZ: pCd1? | 720575940618057095 | 22 |  | Jefferis Lab: A. Javier (1), Irene Salgarella (1). Murthy and Seung Labs: Austin T Burke (23) |
| 57 | SMP & SEZ: pCd1? | 720575940604332460 | 12 |  | Seung Lab: Zhihao Zheng (1). Jefferis and Wilson Labs: Laia Serratos Capdevila (1). Jefferis: A. Javier (1), Bhargavi Parmar (1). Murthy and Seung Labs: Austin T Burke (13). |
| 58 | SMP & SEZ: ADM09p | 720575940642910152 | 12 |  | Scott Lab: Zepeng Yao (6). Murthy and Seung Labs: Austin T Burke (3), Shaina Mae Monungolh (1). Jefferis Lab: Yijie Yin (4), Marina Gkantia (10). |
| 59 | SMP & SEZ: DM3 | 720575940626005330 | 11 |  | Janelia tracers: Tansy Yang (5). Jefferis Lab: Yijie Yin (3). Murthy and Seung Labs: Austin T Burke (7). |
| 60 | SMP & SEZ: Dh44 | 720575940618579505 | 10 |  | Jefferis Lab: Imaan Tamimi (1), A. Javier (4), Rashmita Rana (1). Murthy and Seung Labs: Austin T Burke (13), remer tancontian (7). |
| 61 | SMP & SEZ: pCd1? | 720575940645882420 | 10 |  | Jefferis and Waddell Labs: Joseph Hsu (1). Murthy and Seung Labs: Austin T Burke (18), Joshua Bañez (1). |
| 62 | SMP & SEZ: pCd1? | 720575940638671219 | 10 |  | Jefferis Lab: A. Javier (3), Irene Salgarella (2), Tomke S (2), Varun Sane (5). Murthy and Seung Labs: Claire McKellar (2), remer tancontian (5), Nash Hadjerol (7). |
| 63 | SMP & SEZ: pCd1? | 720575940622119861 | 8 |  | Jefferis Lab: Irene Salgarella (2), Varun Sane (5), Yijie Yin (2). Murthy and Seung Labs: Austin T Burke (21). |
| 64 | SMP & SEZ: DM2_dorsal | 720575940629754588 | 8 |  | Anderson Lab: Altyn Rymbek (1). Jefferis Lab: Yijie Yin (11), Varun Sane (4). Murthy and Seung Labs: Austin T Burke (1), Joshua Bañez (1). |
| 65 | SMP & SEZ: SMPpv2 | 720575940621362044 | 7 |  | Jefferis and Wilson Labs: Laia Serratos Capdevila (1). Jefferis Lab: Yijie Yin (34), Irene Salgarella (1), Imaan Tamimi (7), Chitra Nair (1). Murthy and Seung Labs: Shirleyjoy Serona (1), Austin T Burke (1). |
| 66 | SMP & SEZ: FLA_R.NO_OUT.6 | 720575940623184567 | 7 |  | Jefferis Lab: Katharina Eichler (1), Philipp Schlegel (1). Murthy and Seung Labs: Austin T Burke (1), Ben Silverman (2). |
| 67 | SMP & SEZ: pCd1? | 720575940638719907 | 7 |  | Jefferis and Waddell Labs: Joseph Hsu (1). Murthy and Seung Labs: Austin T Burke (1), Kyle Patrick Willie (6). Wes Murfin (29). |
| 68 | SMP & SEZ: pCd1? | 720575940612572054 | 6 |  | Seung Lab: Zhihao Zheng (1). Jefferis Lab: Irene Salgarella (6). Murthy and Seung Labs: Austin T Burke (37). |
| 69 | SMP & SEZ: DM2 | 720575940608174894 | 5 |  | Anderson Lab: Altyn Rymbek (1), Jefferis and Waddell Labs: Joseph Hsu (14). |
| 70 | SMP & SEZ: DM2 | 720575940635196334 | 5 |  | Jefferis Lab: Irene Salgarella (6). Janelia tracers: Tansy Yang (11). Murthy and Seung Labs: Austin T Burke (7). |
| 71 | SMP & SEZ: pMP5/DM2 | 720575940628732610 | 5 |  | Murthy and Seung Labs: Austin T Burke (4), remer tancontian (5). Jefferis Lab: Irene Salgarella (3), Yijie Yin (22). Janelia tracers: Tansy Yang (2). |
| 72 | SMP & SEZ: SMP_R.FLA_L.29 | 720575940636113264 | 5 |  | Jefferis Lab: Greg Jefferis (1), Laia Serratos (11), Anjali Pandey (1). Dickson Lab: Alisa Poh (1). Murthy and Seung Labs: Nash Hadjerol (11). |
| 73 | SMP & SEZ: pCd1? | 720575940638285184 | 5 | 203 | Murthy and Seung Labs: Austin T Burke (3). Wes Murfin (9). Jefferis Lab: Laia Serratos (4), Varun Sane (4). |
| 74 | Antler L | 720575940622998967 | 54 |  | Murthy and Seung Labs: Austin T Burke (17), Zairene Lenizo (4), Darrel Jay Akiatan (1), Shirleyjoy Serona (12), Shaina Mae Monungolh (2). Jefferis Lab: Yijie Yin (3). |
| 75 | Antler R | 720575940631226439 | 50 | 104 | Murthy and Seung Labs: Austin T Burke (17), Rey Adrian Candilada (3). Jefferis Lab: Varun Sane (7), Yijie Yin (5). |
| 76 | visual projection: SMP_R.SMP_R.201 | 720575940616608837 | 23 |  | Jefferis Lab: Irene Salgarella (1), Rashmita Rana (1), Griffin Badalemente (4). Murthy and Seung Labs: J. Anthony Ocho (12), J. Dolorosa (1). |
| 77 | visual projection: SMP_R.SMP_R.698 | 720575940630512711 | 17 |  | Jefferis Lab: Arti Yadav (1). Murthy and Seung Labs: Doug Bland (19). |
| 78 | visual projection: SMP_L.SMP_L.880 | 720575940636320063 | 15 |  | Murthy and Seung Labs: J. Dolorosa (26). Jefferis Lab: Rashmita Rana (1). |
| 79 | visual projection: SMP_L.SMP_L.362 | 720575940620022960 | 12 | 67 | Kim Lab: Dustin Garner (2). Jefferis Lab: Yijie Yin (16). Murthy and Seung Labs: Ben Silverman (5), remer tancontian (1). |
| 80 | Fan shaped body: DL1_dorsal; left | 720575940626612254 | 29 |  | Jefferis Lab: Rashmita Rana (1), Zeba Vohra (1). Murthy and Seung Labs: Austin T Burke (3), Kyle Patrick Willie (1). |
| 81 | Fan shaped body: FBI1-2 | 720575940642476448 | 17 |  | Murthy and Seung Labs: Jay Gager (1), James Hebditch (4), Kendrick Joules Vinson (1). Jefferis Lab: Varun Sane (6), Yijie Yin (3). |
| 82 | Fan shaped body:<br>FB.FB.473 | 720575940619615936 | 7 |  | Jefferis Lab: Imaan Tamimi (1), A. Javier (1), Laia Serratos (3), Arti Yadav (1), Sangeeta Sisodiya (7). |
| 83 | Fan shaped body:<br>DL1 | 720575940613052200 | 5 | 58 | Jefferis Lab: Varun Sane (3), Griffin Badalemente (1). Murthy and Seung Labs: Shaina Mae Monungolh (1). |
| 84 | Gallinule | 720575940617054621 | 21 |  | Jefferis Lab: Anjali Pandey (1). Murthy and Seung Labs: Mendell Lopez (1), Joshua Bañez (2). |
| 85 | Gallinule | 720575940630349905 | 18 |  | Seeds Hampel Lab: Lucia Kmecova (1). Huetteroth Lab: Wolf Huetteroth (3). Jefferis Lab: Laia Serratos (3). |
| 86 | Gallinule | 720575940629913130 | 7 |  | Murthy and Seung Labs: Zairene Lenizo (12). |
| 87 | Gallinule | 720575940620228449 | 5 |  | Jefferis Lab: Yijie Yin (1). Murthy and Seung Labs: remer tancontian (10), Zairene Lenizo (1). Itisha Joshi (4). |
| 88 | Gallinule | 720575940633711028 | 5 | 56 | Murthy and Seung Labs: Claire McKellar (6) |
| 89 | SEZ:<br>PRW.PRW.213 | 720575940625172016 | 16 |  | Jefferis and Waddell Labs: Joseph Hsu (14). Murthy and Seung Labs: Darrel Jay Akiatan (31), Doug Bland (23), Austin T Burke (1), Ryan Willie (3), remer tancontian (1). |
| 90 | SEZ: PRW.GNG.43 | 720575940630664556 | 8 |  | Jefferis Lab: Marta Costa (1), Sangeeta Sisodiya (2). Seeds Hampel Lab: Alexis E Santana Cruz (2). Murthy and Seung Labs: Zairene Lenizo (3), Darrel Jay Akiatan (5), Ariel Dagohoy (1), Rey Adrian Candilada (2), Joshua Bañez (1), Nash Hadjerol (6), remer tancontian (1). |
| 91 | SEZ:<br>PRW.PRW.312 | 720575940632055521 | 7 |  | Murthy and Seung Labs: J. Dolorosa (3), Shirleyjoy Serona (103). Jefferis Lab: Arti Yadav (1). |
| 92 | SEZ: PRW.PRW.82 | 720575940616860758 | 6 |  | Jefferis Lab: Arti Yadav (1). Murthy and Seung Labs: Joshua Bañez (7), Zairene Lenizo (3), Doug Bland (12), J. Dolorosa (2), remer tancontian (1). |
| 93 | SEZ:<br>PRW.PRW.164 | 720575940622457579 | 6 | 43 | Jefferis Lab: Arti Yadav (1). Murthy and Seung Labs: Michelle Pantujan (2), Shirleyjoy Serona (3). |
|  |  |  |  | 1742 |  |

**Table S3:**
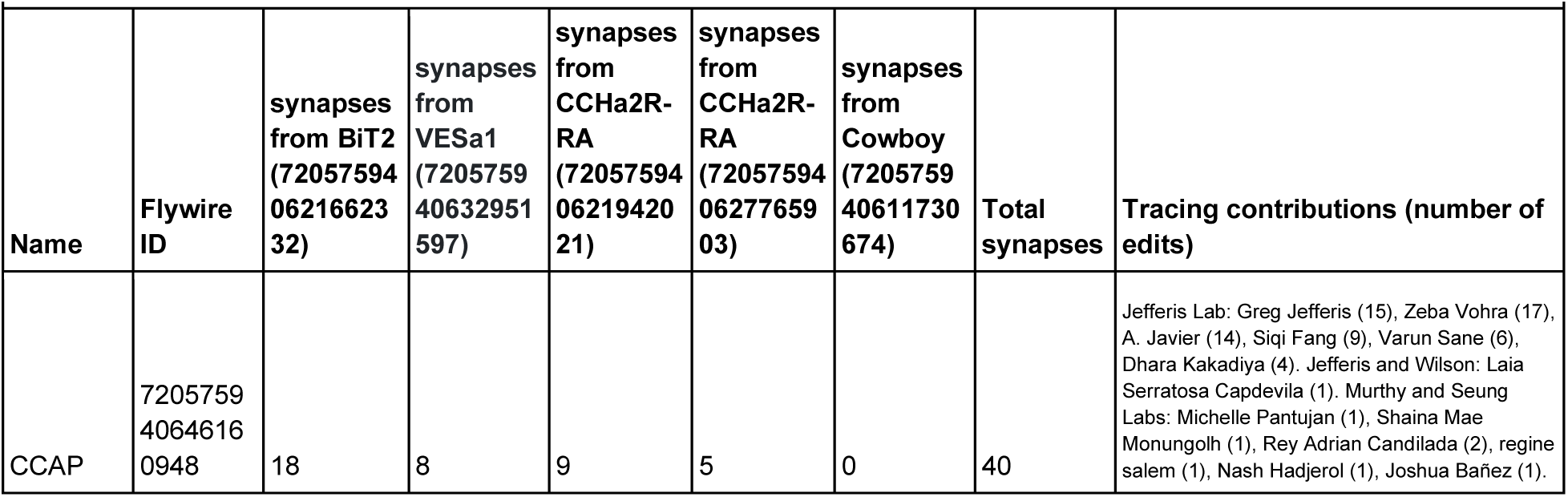
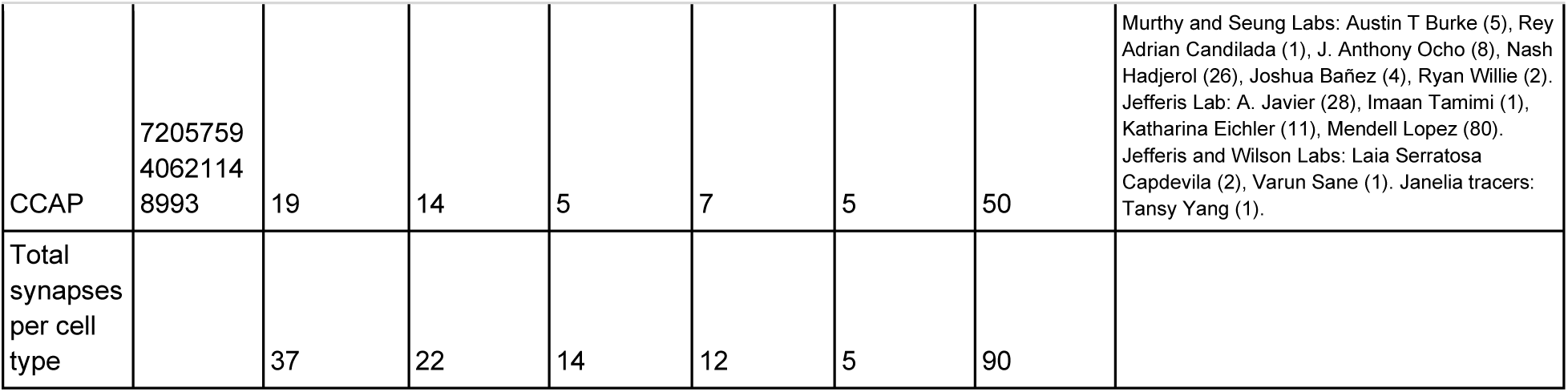
CCAP presynaptic partners.

| Table S3: CCAP presynaptic partners |  |  |  |  |  |  |  |  |
| --- | --- | --- | --- | --- | --- | --- | --- | --- |
| Name | Flywire ID | synapses from BiT2 (72057594 06216623 32) | synapses from VESa1 (7205759 40632951 597) | synapses from CCHa2R-RA (72057594 06219420 21) | synapses from CCHa2R-RA (72057594 06277659 03) | synapses from Cowboy (7205759 40611730 674) | Total synapses | Tracing contributions (number of edits) |
| CCAP | 7205759 4064616 0948 | 18 | 8 | 9 | 5 | 0 | 40 | Jefferis Lab: Greg Jefferis (15), Zeba Vohra (17), A. Javier (14), Siqi Fang (9), Varun Sane (6), Dhara Kakadiya (4). Jefferis and Wilson: Laia Serratos Capdevila (1). Murthy and Seung Labs: Michelle Pantujan (1), Shaina Mae Monungolh (1), Rey Adrian Candilada (2), regine salem (1), Nash Hadjerol (1), Joshua Bañez (1). |
| CCAP | 7205759<br>4062114<br>8993 | 19 | 14 | 5 | 7 | 5 | 50 | Murthy and Seung Labs: Austin T Burke (5), Rey Adrian Candilada (1), J. Anthony Ocho (8), Nash Hadjerol (26), Joshua Bañez (4), Ryan Willie (2).<br>Jefferis Lab: A. Javier (28), Imaan Tamimi (1), Katharina Eichler (11), Mendell Lopez (80).<br>Jefferis and Wilson Labs: Laia Serratosa Capdevila (2), Varun Sane (1). Janelia tracers: Tansy Yang (1). |
| Total synapses per cell type |  | 37 | 22 | 14 | 12 | 5 | 90 |  |

**Table S4:**
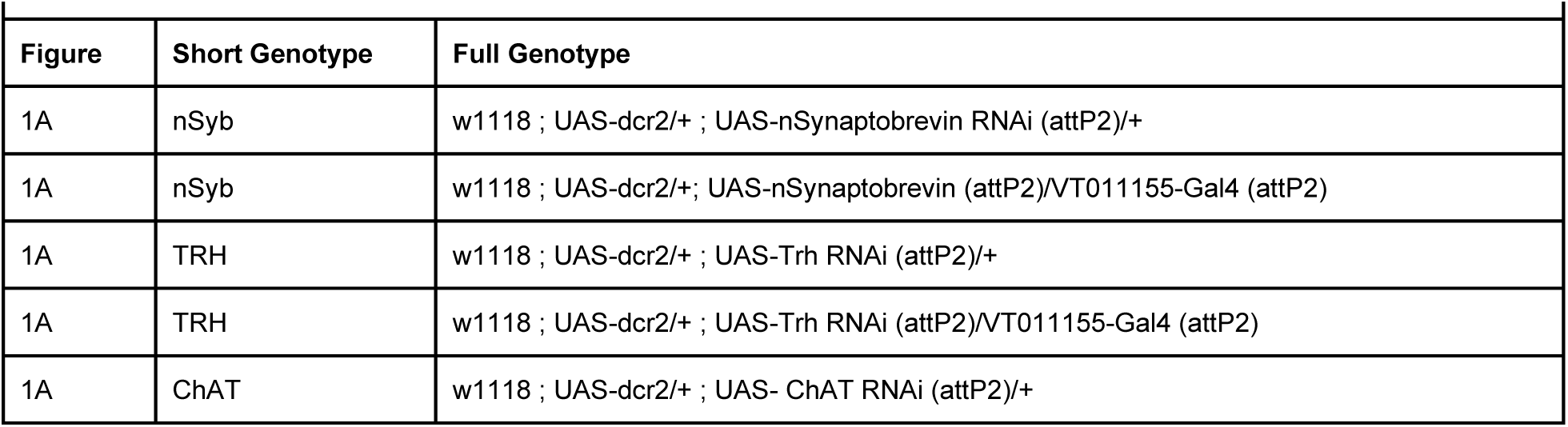
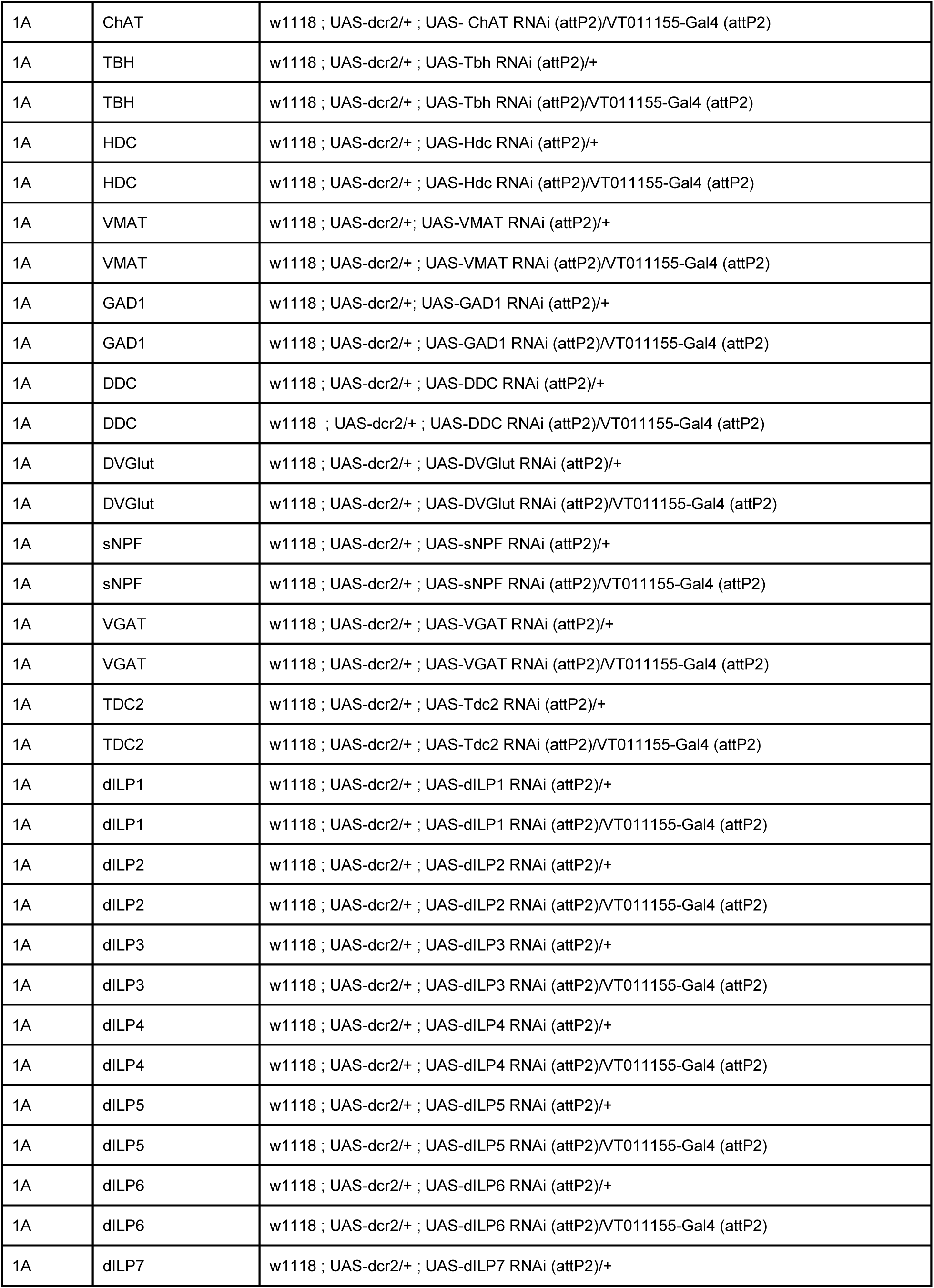
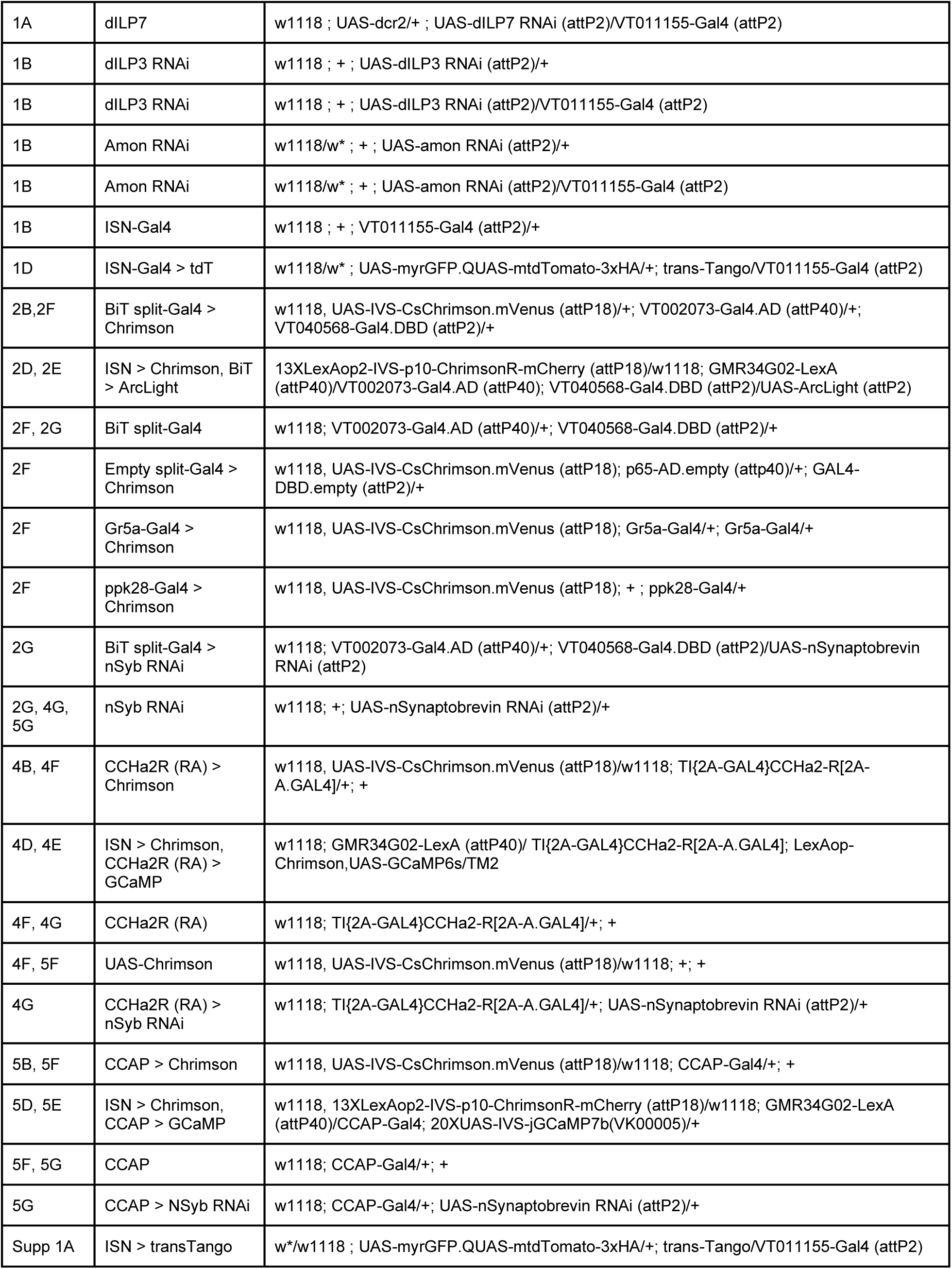
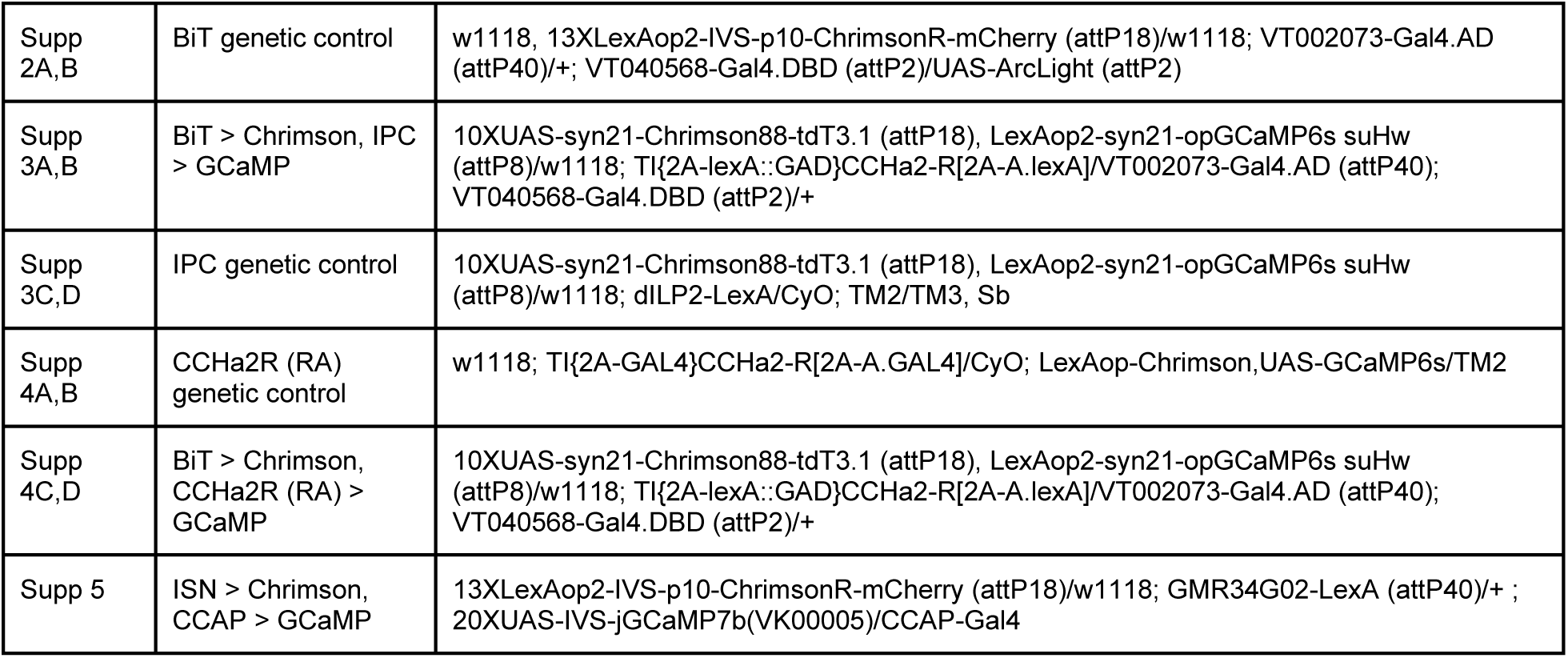
Fly genotypes in figures.

**Figure 1 - supplement 1.**
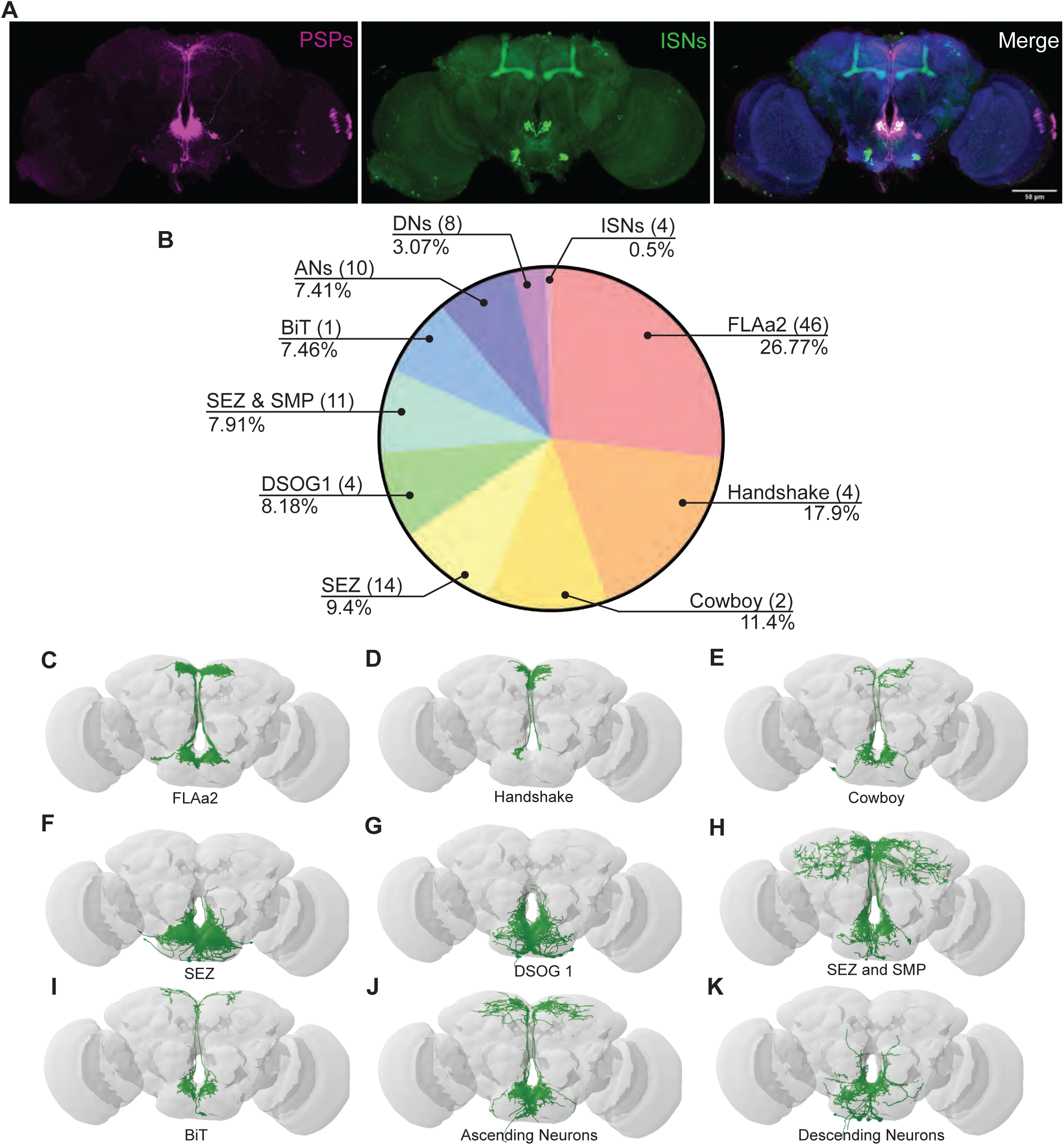
ISN postsynaptic partners labeled by trans-Tango and EM. (A) Expression of trans-Tango ligand in the ISNs (green) and postsynaptic partners (PSPs) (magenta). nc82 staining in blue. (B) Distribution of synaptic output from the ISNs divided by cell class or brain region. Total of 4050 synapses from the ISNs and 104 postsynaptic partners. FLAa2 (46 neurons) receive 26.77% of all ISN output, Handshake (4 neurons) 17.9%, Cowboy (2 neurons) 11.4%, neurons located in the subesophageal zone (SEZ) (14 neurons) 9.04%, DSOG1 (4 neurons) 8.18%, neurons with neurites in the subesophageal zone and superior medial protocerebrum (SEZ & SMP) (11 neurons) 7.91%, BiT (1 neuron) 7.46%, Ascending neurons (ANs) (10 neurons) 4.41%, Descending neurons (DNs) (8 neurons) 3.07%, and ISNs (4 neurons) 0.5%. Only postsynaptic partners with 5 or more synapses were considered for this analysis. Reconstruction of FLAa2 neurons (C), Handshake neurons (D), Cowboy neurons (E), neurons innervating the SEZ (F), DSOG1 neurons (G), neurons innervating the SEZ & SMP (H), BiT (I), ascending neurons (J), descending neurons (K).

**Figure 2 - supplement 2.**
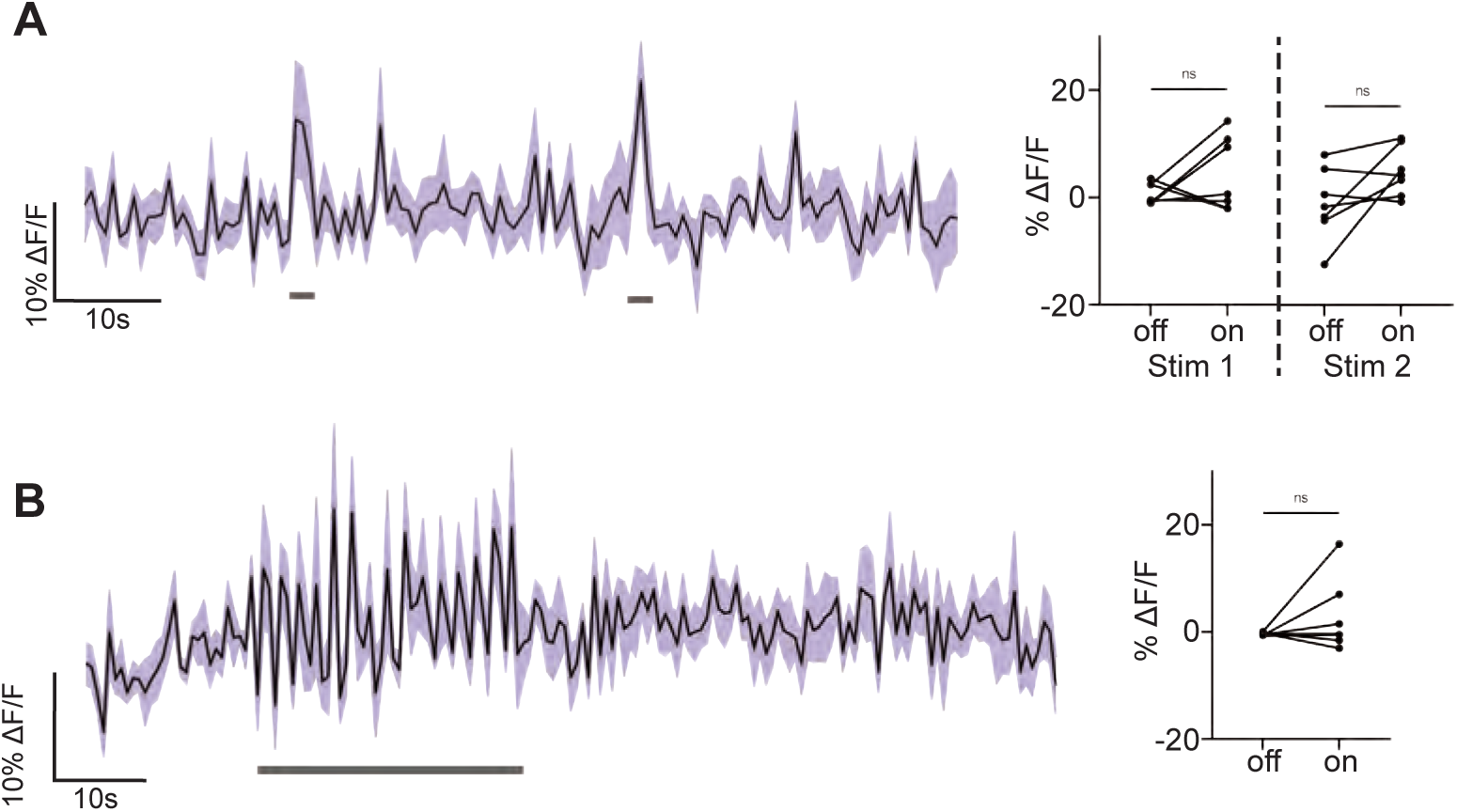
BiT genetic control functional imaging. Genetic controls: we expressed the light sensitive ion channel Chrimson without the ISN-LexA driver and exposed the brains to 660nm LED. We expressed the voltage sensor ArcLight in BiT and imaged it with a 2 photon microscope. (A) ArcLight response of BiT soma to 2s LED exposure or (B) 30s LED exposure. Left: Scatter plot shows mean +/- SEM of all flies imaged, gray bars represent LED stimulation. Right: Quantification of mean fluorescence intensity before stim (off) and during stim (on), each dot represents one fly. Paired Wilcoxon or paired t-test. n=7 flies.

**Figure 2 - supplement 2.**
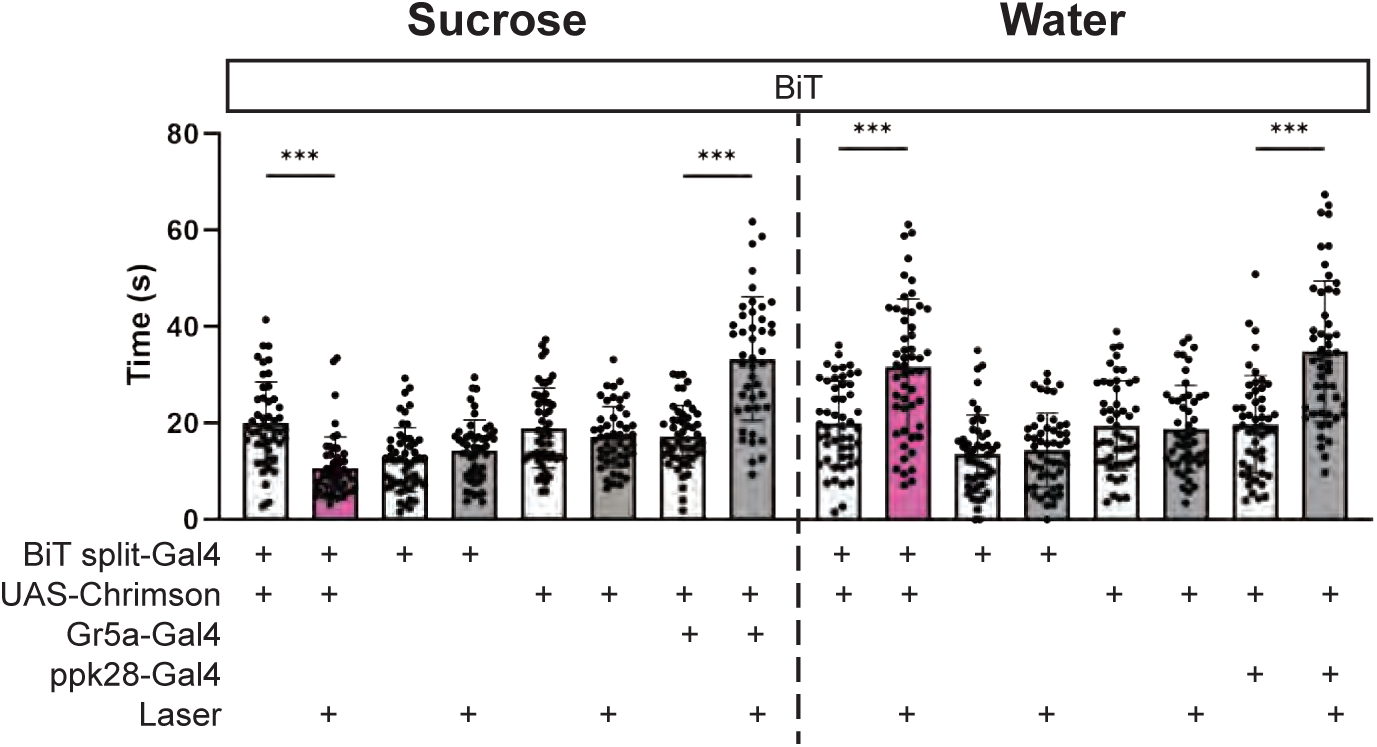
BiT optogenetic activation inges-tion phenotype. Temporal consumption assay for 1M sucrose or water during acute optogenetic activation of BiT with Chrimson. Represented are the mean, and the 10-90 percentile. Data was analyzed using Kruskal-Wallis test, followed by multiple compari-sons against the no laser control. n=44-54 animals/geno-type. ***p<0.001

**Figure 3 - supplement 1.**
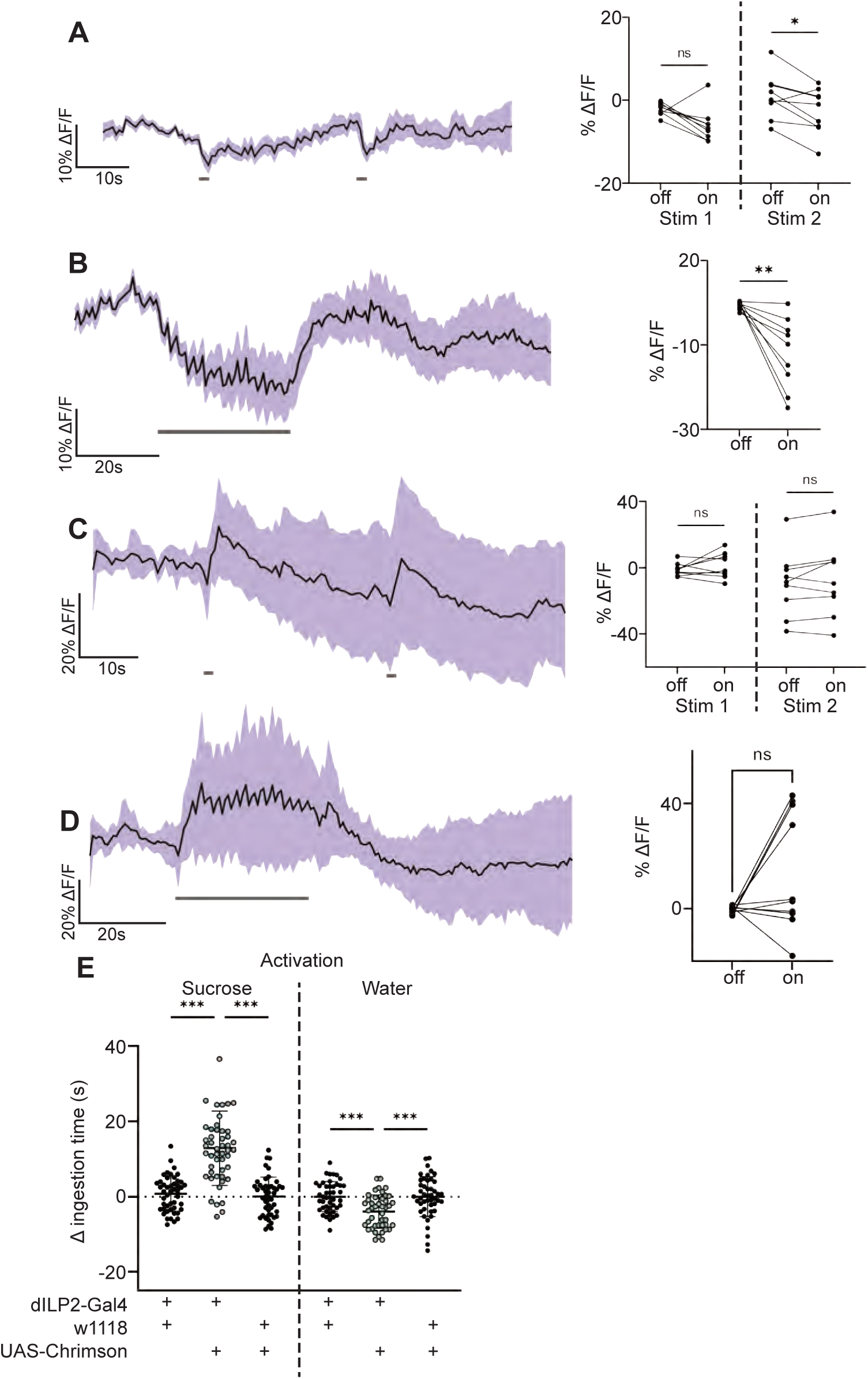
IPC response to BiT stimulation. A and B: We expressed the light sensitive ion channel Chrimson in BiT and optogenetically stimulated it with 660nm LED. We expressed the calcium sensor GCaMP in the IPCs and imaged them with a 2 photon microscope. (A) Calcium response of IPC somas to 2s optogenetic activation of BiT or (B) 30s optogenetic activation of BiT. Left: Scatter plot shows mean +/- SEM of all flies imaged, gray bars represent LED stimulation. Right: Quantification of mean fluores-cence intensity before stim (off) and during stim (on), each dot represents one fly. Paired Wilcoxon test or paired t-test. n=9 flies. C and D: Genetic controls: we expressed the light sensitive ion channel Chrimson without the BiT-split Gal4 driver and exposed the brains to 660nm LED. We expressed the calcium sensor GCaMP in the IPCs and imaged them with a 2 photon microscope. (C) Calcium response of IPC somas to 2s LED exposure or (D) 30s LED exposure. Left: Scatter plot shows mean +/- SEM of all flies imaged, gray bars represent LED stimulation. Right: Quantifica-tion of mean fluorescence intensity before stim (off) and during stim (on), each dot represents one fly. Paired t-test. n=9-10 flies. (E) Temporal consumption assay for 1M sucrose or water during acute optogenetic activation of IPCs with Chrimson. Ingestion time of females exposed to light normal-ized to dark controls of indicated genotype. One-way ANOVA with Holm-Šídák multiple comparison test. n=43-49 animals/genotype. *p<0.05, **p<0.01, ***p<0.001

**Figure 3 - supplement 2.**
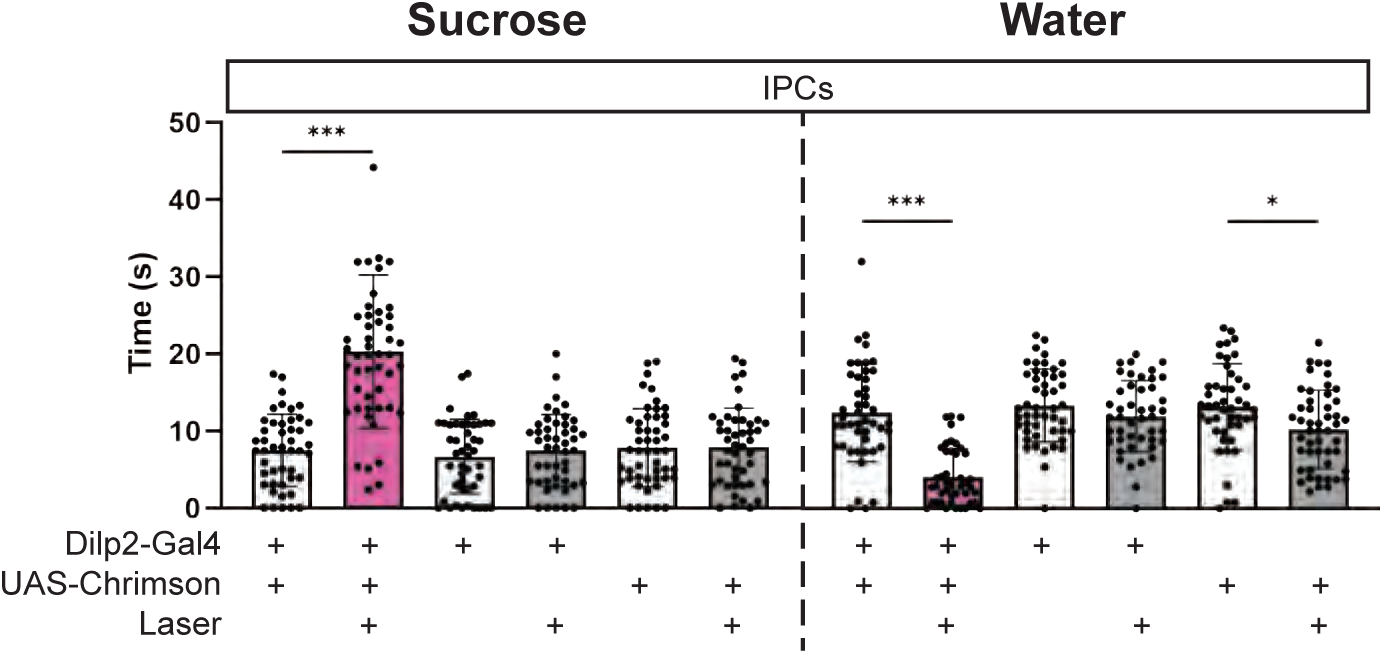
IPC optogenetic activation ingestion phenotype. Temporal consumption assay for 1M sucrose or water during acute optogenetic activation of IPCs with Chrimson. Represented are the mean, and the 10-90 percentile. Data was analyzed using One-way ANOVA, followed by multiple comparisons against the no laser control. n=43-49 animals/genotype. *p<0.05, ***p<0.001

**Figure 4 - supplement 1.**
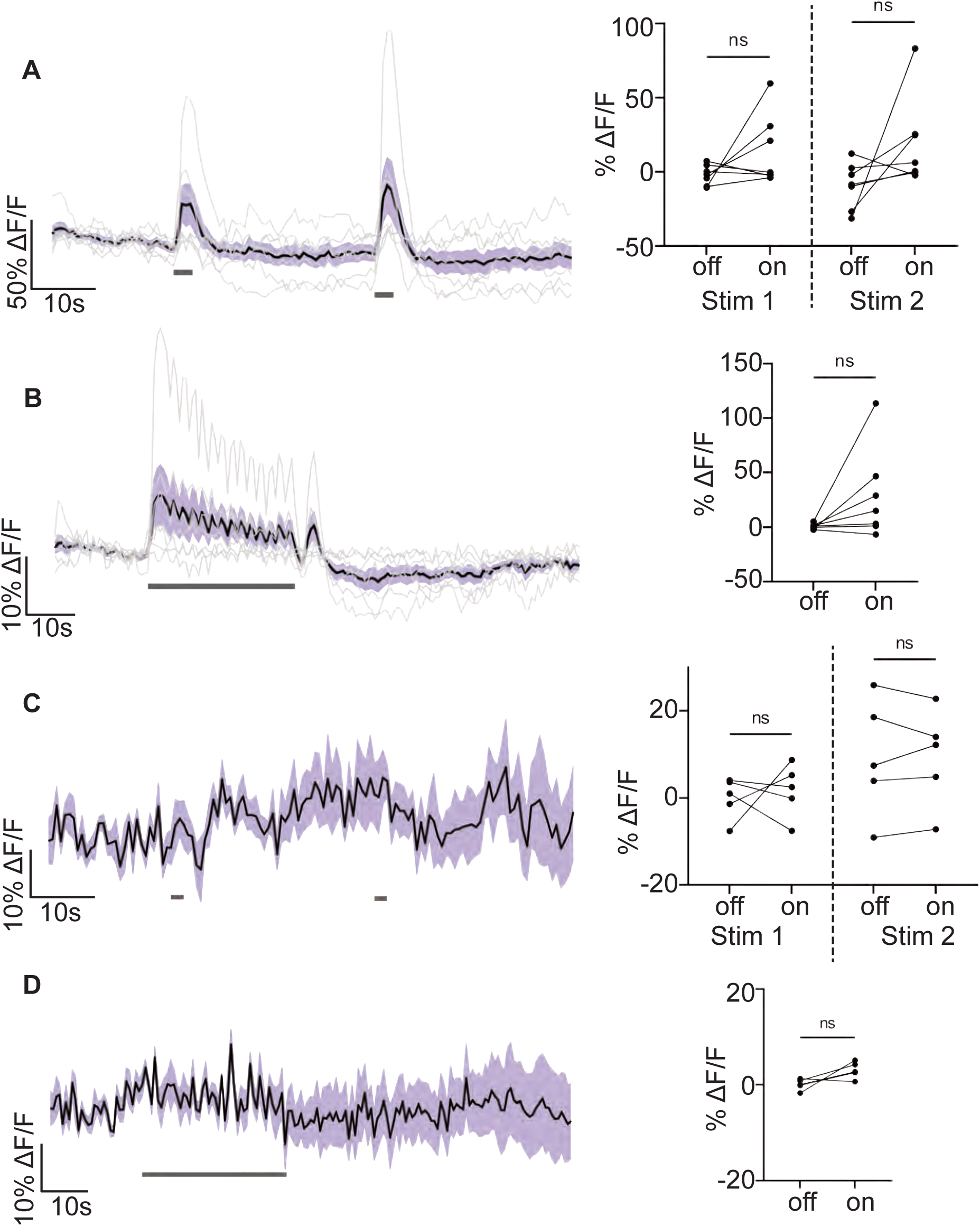
CCHa2R-RA genetic controls functional imaging and response to BiT optogenetic stimulation. A and B: Genetic controls: we expressed the light sensitive ion channel Chrimson without the ISN-LexA driver and exposed the brains to 660nm LED. We expressed the calcium sensor GCaMP in the CCHa2R-RA neurons and imaged them with a 2 photon microscope. (A) Calcium response of CCHa2R-RA SEZ neurites to 2s LED exposure or (B) 30s LED exposure. Left: Scatter plot shows mean +/- SEM of all flies imaged, with individual traces in gray, gray bars represent LED stimulation. Right: Quantification of mean fluorescence intensity before stim (off) and during stim (on), each dot represents one fly. Paired Wilcoxon test and paired t-test. n=7 flies. C and D: We expressed the light sensitive ion channel Chrimson in BiT and optogenetically stimulated it with 660nm LED. We expressed the calcium sensor GCaMP in the CCHa2R-RA neurons and imaged them with a 2 photon microscope. (C) Calcium response of CCHa2R-RA somas to 2s optogenetic stimulation of BiT or (D) 30s optogenetic stimulation of BiT. Left: Scatter plot shows mean +/- SEM of all flies imaged, gray bars represent LED stimulation. Right: Quantification of mean fluores-cence intensity before stim (off) and during stim (on), each dot represents one fly. Paired t-test. n=5 flies.

**Figure 4 - supplement 2.**
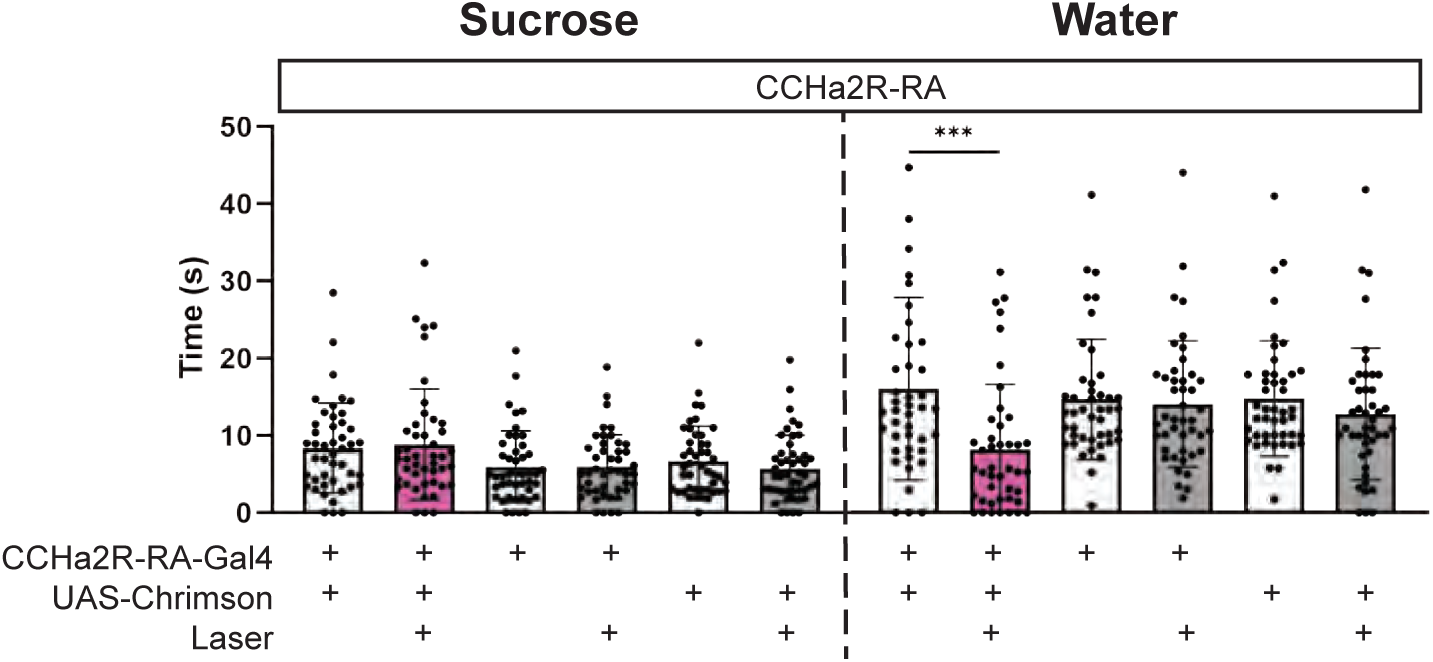
CCHa2R-RA optogenetic activation ingestion phenotype. Temporal consumption assay for 1M sucrose or water during acute optogenetic activation of CCHa2R-RA neurons with Chrimson. Represented are the mean, and the 10-90 percentile. Data was analyzed using Krus-kal-Wallis test, followed by multiple comparisons against the no laser control. n=42-47 animals/genotype. ***p<0.001

**Figure 5 - supplement 1.**
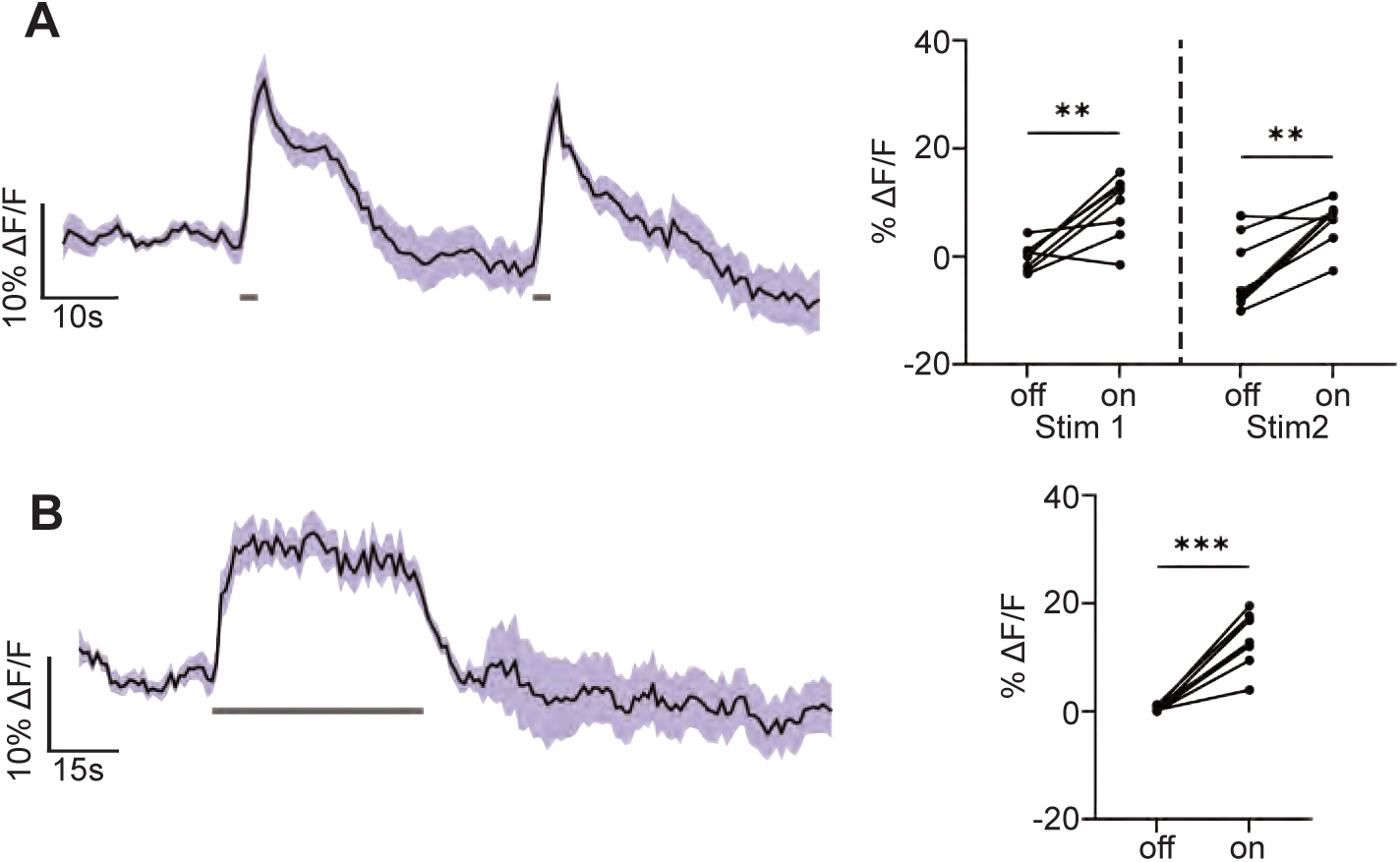
CCAP response to ISN activation using another CCAP-Gal4 driver. We expressed the light sensitive ion channel Chrimson in the ISNs and optoge-netically stimulated them with 660nm LED. We expressed the calcium sensor GCaMP in the CCAP neurons and imaged them with a 2 photon microscope. (A) Calcium response of CCAP neurites to 2s optogenetic stimulation of the ISNs or (B) 30s optogenetic stimulation of the ISNs. Left: Scatter plot shows mean +/- SEM of all flies imaged, gray bars represent LED stimulation. Right: Quantification of mean fluorescence intensity before stim (off) and during stim (on), each dot represents one fly. Paired t-test. n=8 flies. **p<0.01, ***p<0.001

**Figure 5 - supplement 2.**
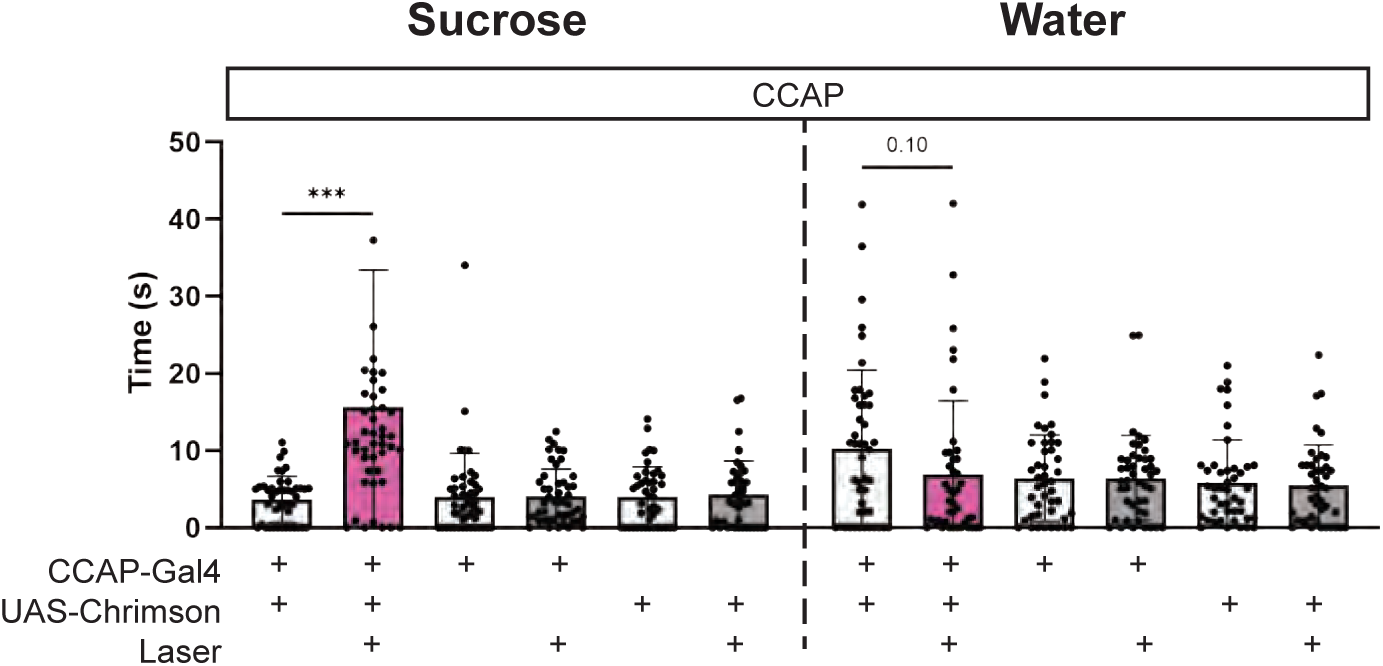
CCAP optogenetic activa-tion ingestion phenotype. Temporal consumption assay for 1M sucrose or water during acute optogenetic activation of CCAP neurons with Chrimson. Represented are the mean, and the 10-90 percentile. Data was analyzed using Kruskal-Wallis test, followed by multiple compari-sons against the no laser control. n=42-48 animals/gen-otype. ***p<0.001

## Notes

### Competing Interest Statement

The authors have declared no competing interest.

### Summary of Updates

revised manuscript to address elife review

## REFERENCES

Aponte, Y., Atasoy, D., & Sternson, S. M. (2011). AGRP neurons are sufficient to orchestrate feeding behavior rapidly and without training. Nat Neurosci, 14(3), 351–355. https://doi.org/10.1038/nn.2739

Badisco, L., Van Wielendaele, P., & Vanden Broeck, J. (2013). Eat to reproduce: a key role for the insulin signaling pathway in adult insects. Front Physiol, 4, 202. https://doi.org/10.3389/fphys.2013.00202

Baker, C. A., McKellar, C., Pang, R., Nern, A., Dorkenwald, S., Pacheco, D. A., Eckstein, N., Funke, J., Dickson, B. J., & Murthy, M. (2022). Neural network organization for courtship-song feature detection in Drosophila. Curr Biol, 32(15), 3317–3333 e3317. https://doi.org/10.1016/j.cub.2022.06.019

Bates, A. S., Manton, J. D., Jagannathan, S. R., Costa, M., Schlegel, P., Rohlfing, T., & Jefferis, G. S. (2020). The natverse, a versatile toolbox for combining and analysing neuroanatomical data. Elife, 9. https://doi.org/10.7554/eLife.53350

Biglou, S. G., Bendena, W. G., & Chin-Sang, I. (2021). An overview of the insulin signaling pathway in model organisms Drosophila melanogaster and Caenorhabditis elegans. Peptides, 145, 170640. https://doi.org/10.1016/j.peptides.2021.170640

Brogiolo, W., Stocker, H., Ikeya, T., Rintelen, F., Fernandez, R., & Hafen, E. (2001). An evolutionarily conserved function of the Drosophila insulin receptor and insulin-like peptides in growth control. Current biology, 11(4), 213–221.

Buhmann, J., Sheridan, A., Malin-Mayor, C., Schlegel, P., Gerhard, S., Kazimiers, T., Krause, R., Nguyen, T. M., Heinrich, L., Lee, W. A., Wilson, R., Saalfeld, S., Jefferis, G., Bock, D. D., Turaga, S. C., Cook, M., & Funke, J. (2021). Automatic detection of synaptic partners in a whole-brain Drosophila electron microscopy data set. Nat Methods, 18(7), 771–774. https://doi.org/10.1038/s41592-021-01183-7

Burnett, C. J., Li, C., Webber, E., Tsaousidou, E., Xue, S. Y., Bruning, J. C., & Krashes, M. J. (2016). Hunger-Driven Motivational State Competition. Neuron, 92(1), 187–201. https://doi.org/10.1016/j.neuron.2016.08.032

Cannell, E., Dornan, A. J., Halberg, K. A., Terhzaz, S., Dow, J. A. T., & Davies, S. A. (2016). The corticotropin-releasing factor-like diuretic hormone 44 (DH44) and kinin neuropeptides modulate desiccation and starvation tolerance in Drosophila melanogaster. Peptides, 80, 96–107. https://doi.org/10.1016/j.peptides.2016.02.004

Chen, C., Jack, J., & Garofalo, R. S. (1996). The Drosophila insulin receptor is required for normal growth. Endocrinology, 137(3), 846–856.

Chen, T. W., Wardill, T. J., Sun, Y., Pulver, S. R., Renninger, S. L., Baohan, A., Schreiter, E. R., Kerr, R. A., Orger, M. B., Jayaraman, V., Looger, L. L., Svoboda, K., & Kim, D. S. (2013). Ultrasensitive fluorescent proteins for imaging neuronal activity. Nature, 499(7458), 295–300. https://doi.org/10.1038/nature12354

Claeys, I., Simonet, G., Poels, J., Van Loy, T., Vercammen, L., De Loof, A., & Broeck, J. V. (2002). Insulin-related peptides and their conserved signal transduction pathway. Peptides, 23(4), 807–816.

Clancy, D. J., Gems, D., Harshman, L. G., Oldham, S., Stocker, H., Hafen, E., Leevers, S. J., & Partridge., L. (2001). Extension of Life-Span by Loss of CHICO, a Drosophila Insulin Receptor Substrate Protein. Science, 292(5514), 104–106.

Deng, B., Li, Q., Liu, X., Cao, Y., Li, B., Qian, Y., Xu, R., Mao, R., Zhou, E., Zhang, W., Huang, J., & Rao, Y. (2019). Chemoconnectomics: Mapping Chemical Transmission in Drosophila. Neuron, 101(5), 876–893 e874. https://doi.org/10.1016/j.neuron.2019.01.045

Dethier, V. G. (1976). The hungry fly: A physiological study of the behavior associated with feeding.

Dionne, H., Hibbard, K. L., Cavallaro, A., Kao, J. C., & Rubin, G. M. (2018). Genetic Reagents for Making Split-GAL4 Lines in Drosophila. Genetics, 209(1), 31–35. https://doi.org/10.1534/genetics.118.300682

Dorkenwald, S., McKellar, C. E., Macrina, T., Kemnitz, N., Lee, K., Lu, R., Wu, J., Popovych, S., Mitchell, E., & Nehoran, B. (2022). FlyWire: online community for whole-brain connectomics. Nature methods, 19(1), 119–128.

Eckstein, N., Bates, A. S., Du, M., Hartenstein, V., Jefferis, G. S. X. E., & Funke, J. (2020). Neurotransmitter Classification from Electron

Microscopy Images at Synaptic Sites in Drosophila. bioRxiv. https://doi.org/10.1101/2020.06.12.148775

Eiselt, A. K., Chen, S., Chen, J., Arnold, J., Kim, T., Pachitariu, M., & Sternson, S. M. (2021). Hunger or thirst state uncertainty is resolved by outcome evaluation in medial prefrontal cortex to guide decision-making. Nat Neurosci, 24(7), 907–912. https://doi.org/10.1038/s41593-021-00850-4

Fernandez, R., Tabarini, D., Azpiazu, N., Frasch, M., & Schlessinger, J. (1995). The Drosophila insulin receptor homolog: a gene essential for embryonic development encodes two receptor isoforms with different signaling potential. EMBO J, 14(14), 3373–3384. https://doi.org/10.1002/j.1460-2075.1995.tb07343.x

Gáliková, M., Diesner, M., Klepsatel, P., Hehlert, P., Xu, Y., Bickmeyer, I., Predel, R., & Kuhnlein, R. P. (2015). Energy Homeostasis Control in Drosophila Adipokinetic Hormone Mutants. Genetics, 201(2), 665–683. https://doi.org/10.1534/genetics.115.178897

Gáliková, M., Dircksen, H., & Nässel, D. R. (2018). The thirsty fly: Ion transport peptide (ITP) is a novel endocrine regulator of water homeostasis in Drosophila. PLoS Genet, 14(8), e1007618. https://doi.org/10.1371/journal.pgen.1007618

Garofalo, R. S. (2002). Genetic analysis of insulin signaling in Drosophila. Trends in Endocrinology & Metabolism, 13(4), 156–162.

Gizowski, C., & Bourque, C. W. (2018). The neural basis of homeostatic and anticipatory thirst. Nat Rev Nephrol, 14(1), 11–25. https://doi.org/10.1038/nrneph.2017.149

Gordon, M. D., & Scott, K. (2009). Motor control in a Drosophila taste circuit. Neuron, 61(3), 373–384. https://doi.org/10.1016/j.neuron.2008.12.033

Graham, M., Shutter, J. R., Sarmiento, U., Sarosi, l., & Stark, K. L. (1997). Overexpression of Agrt leads to obesity in transgenic mice. Nature genetics, 17(3), 273–274.

Grönke, S., Clarke, D. F., Broughton, S., Andrews, T. D., & Partridge, L. (2010). Molecular evolution and functional characterization of Drosophila insulin-like peptides. PLoS Genet, 6(2), e1000857. https://doi.org/10.1371/journal.pgen.1000857

Hartenstein, V. (2006). The neuroendocrine system of invertebrates: a developmental and evolutionary perspective. J Endocrinol, 190(3), 555–570. https://doi.org/10.1677/joe.1.06964

Heinrich, L., Funke, J., Pape, C., Nunez-Iglesias, J., & Saalfeld, S. (2018). Synaptic cleft segmentation in non-isotropic volume electron microscopy of the complete drosophila brain. Medical Image Computing and Computer Assisted Intervention–MICCAI 2018: 21st International Conference, Granada, Spain, September 16-20, 2018, Proceedings, Part II 11,

Ida, T., Takahashi, T., Tominaga, H., Sato, T., Sano, H., Kume, K., Ozaki, M., Hiraguchi, T., Shiotani, H., Terajima, S., Nakamura, Y., Mori, K., Yoshida, M., Kato, J., Murakami, N., Miyazato, M., Kangawa, K., & Kojima, M. (2012). Isolation of the bioactive peptides CCHamide-1 and CCHamide-2 from Drosophila and their putative role in appetite regulation as ligands for G protein-coupled receptors. Front Endocrinol (Lausanne), 3, 177. https://doi.org/10.3389/fendo.2012.00177

Jin, L., Han, Z., Platisa, J., Wooltorton, J. R., Cohen, L. B., & Pieribone, V. A. (2012). Single action potentials and subthreshold electrical events imaged in neurons with a fluorescent protein voltage probe. Neuron, 75(5), 779–785. https://doi.org/10.1016/j.neuron.2012.06.040

Jourjine, N. (2017). Hunger and thirst interact to regulate ingestive behavior in flies and mammals. BioEssays, 39(5). https://doi.org/10.1002/bies.201600261

Jourjine, N., Mullaney, B. C., Mann, K., & Scott, K. (2016). Coupled Sensing of Hunger and Thirst Signals Balances Sugar and Water Consumption. Cell, 166(4), 855–866. https://doi.org/10.1016/j.cell.2016.06.046

Kim, J., & Neufeld, T. P. (2015). Dietary sugar promotes systemic TOR activation in Drosophila through AKH-dependent selective secretion of Dilp3. Nat Commun, 6, 6846. https://doi.org/10.1038/ncomms7846

Klapoetke, N. C., Murata, Y., Kim, S. S., Pulver, S. R., Birdsey-Benson, A., Cho, Y. K., Morimoto, T. K., Chuong, A. S., Carpenter, E. J., Tian, Z., Wang, J., Xie, Y., Yan, Z., Zhang, Y., Chow, B. Y., Surek, B., Melkonian, M., Jayaraman, V., Constantine-Paton, M., . . . Boyden, E. S. (2014). Independent optical excitation of distinct neural populations. Nat Methods, 11(3), 338–346. https://doi.org/10.1038/nmeth.2836

Landayan, D., Wang, B. P., Zhou, J., & Wolf, F. W. (2021). Thirst interneurons that promote water seeking and limit feeding behavior in Drosophila. Elife, 10. https://doi.org/10.7554/eLife.66286

Laturney, M., Sterne, G. R., & Scott, K. (2022). Mating activates neuroendocrine pathways signaling hunger in Drosophila females. bioRxiv. https://doi.org/10.1101/2022.10.19.512959

Lee, Y. J., Yang, C. P., Miyares, R. L., Huang, Y. F., He, Y., Ren, Q., Chen, H. M., Kawase, T., Ito, M., Otsuna, H., Sugino, K., Aso, Y., Ito, K., & Lee, T. (2020). Conservation and divergence of related neuronal lineages in the Drosophila central brain. Elife, 9. https://doi.org/10.7554/eLife.53518

Li, P. H., Lindsey, L. F., Januszewski, M., Tyka, M., Maitin-Shepard, J., Blakely, T., & Jain, V. (2019). Automated Reconstruction of a Serial-Section EM Drosophila Brain with Flood-Filling Networks and Local Realignment. Microscopy and Microanalysis, 25(S2), 1364–1365. https://doi.org/10.1017/s1431927619007554

Lin, S., Owald, D., Chandra, V., Talbot, C., Huetteroth, W., & Waddell, S. (2014). Neural correlates of water reward in thirsty Drosophila. Nat Neurosci, 17(11), 1536–1542. https://doi.org/10.1038/nn.3827

Liu, W. W., & Wilson, R. I. (2013). Glutamate is an inhibitory neurotransmitter in the Drosophila olfactory system. Proc Natl Acad Sci U S A, 110(25), 10294–10299. https://doi.org/10.1073/pnas.1220560110

Meissner, G. W., Luo, S. D., Dias, B. G., Texada, M. J., & Baker, B. S. (2016). Sex-specific regulation of Lgr3 in Drosophila neurons. Proc Natl Acad Sci U S A, 113(9), E1256–1265. https://doi.org/10.1073/pnas.1600241113

Min, S., Chae, H. S., Jang, Y. H., Choi, S., Lee, S., Jeong, Y. T., Jones, W. D., Moon, S. J., Kim, Y. J., & Chung, J. (2016). Identification of a Peptidergic Pathway Critical to Satiety Responses in Drosophila. Curr Biol, 26(6), 814–820. https://doi.org/10.1016/j.cub.2016.01.029

Nässel, D. R., Kubrak, O. I., Liu, Y., Luo, J., & Lushchak, O. V. (2013). Factors that regulate insulin producing cells and their output in Drosophila. Front Physiol, 4, 252. https://doi.org/10.3389/fphys.2013.00252

Nässel, D. R., Liu, Y., & Luo, J. (2015). Insulin/IGF signaling and its regulation in Drosophila. Gen Comp Endocrinol, 221, 255–266. https://doi.org/10.1016/j.ygcen.2014.11.021

Nässel, D. R., & Vanden Broeck, J. (2016). Insulin/IGF signaling in Drosophila and other insects: factors that regulate production, release and post-release action of the insulin-like peptides. Cell Mol Life Sci, 73(2), 271–290. https://doi.org/10.1007/s00018-015-2063-3

Nässel, D. R., & Zandawala, M. (2020). Hormonal axes in Drosophila: regulation of hormone release and multiplicity of actions. Cell Tissue Res, 382(2), 233–266. https://doi.org/10.1007/s00441-020-03264-z

Nässel, D. R., & Zandawala, M. (2022). Endocrine cybernetics: neuropeptides as molecular switches in behavioural decisions. Open Biol, 12(7), 220174. https://doi.org/10.1098/rsob.220174

Nern, A., Pfeiffer, B. D., & Rubin, G. M. (2015). Optimized tools for multicolor stochastic labeling reveal diverse stereotyped cell arrangements in the fly visual system. Proc Natl Acad Sci U S A, 112(22), E2967–2976. https://doi.org/10.1073/pnas.1506763112

Ohhara, Y., Kobayashi, S., Yamakawa-Kobayashi, K., & Yamanaka, N. (2018). Adult-specific insulin-producing neurons in Drosophila melanogaster. J Comp Neurol, 526(8), 1351–1367. https://doi.org/10.1002/cne.24410

Orchard, I. (1987). Adipokinetic hormones—an update. Journal of insect physiology, 33(7), 451–463.

Otsuna, H., Ito, M., & Kawase, T. (2018). Color depth MIP mask search: a new tool to expedite SplitGAL4 creation. bioRxiv. https://doi.org/10.1101/318006

Palovcik, R. A., Phillips, M. I., Kappy, M. S., & Raizada, M. K. (1984). Insulin inhibits pyramidal neurons in hippocampal slices. Brain research, 309(1), 187–191.

Pfeiffer, B. D., Jenett, A., Hammonds, A. S., Ngo, T.-T. B., Misra, S., Murphy, C., Scully, A., Carlson, J. W., Wan, K. H., & Laverty, T. R. (2008). Tools for neuroanatomy and neurogenetics in Drosophila. Proceedings of the National Academy of Sciences, 105(28), 9715–9720.

Pool, A. H., Kvello, P., Mann, K., Cheung, S. K., Gordon, M. D., Wang, L., & Scott, K. (2014). Four GABAergic interneurons impose feeding restraint in Drosophila. Neuron, 83(1), 164–177. https://doi.org/10.1016/j.neuron.2014.05.006

Qi, W., Wang, G., & Wang, L. (2021). A novel satiety sensor detects circulating glucose and suppresses food consumption via insulin-producing cells in Drosophila. Cell Res, 31(5), 580–588. https://doi.org/10.1038/s41422-020-00449-7

Reiher, W., Shirras, C., Kahnt, J., Baumeister, S., Isaac, R. E., & Wegener, C. (2011). Peptidomics and peptide hormone processing in the Drosophila midgut. J Proteome Res, 10(4), 1881–1892. https://doi.org/10.1021/pr101116g

Ren, G. R., Hauser, F., Rewitz, K. F., Kondo, S., Engelbrecht, A. F., Didriksen, A. K., Schjott, S. R., Sembach, F. E., Li, S., Sogaard, K. C., Sondergaard, L., & Grimmelikhuijzen, C. J. (2015). CCHamide-2 Is an Orexigenic Brain-Gut Peptide in Drosophila. PLoS One, 10(7), e0133017. https://doi.org/10.1371/journal.pone.0133017

Saalfeld, S., Cardona, A., Hartenstein, V., & Tomancak, P. (2009). CATMAID: collaborative annotation toolkit for massive amounts of image data. Bioinformatics, 25(15), 1984–1986. https://doi.org/10.1093/bioinformatics/btp266

Sano, H., Nakamura, A., Texada, M. J., Truman, J. W., Ishimoto, H., Kamikouchi, A., Nibu, Y., Kume, K., Ida, T., & Kojima, M. (2015). The Nutrient-Responsive Hormone CCHamide-2 Controls Growth by Regulating Insulin-like Peptides in the Brain of Drosophila melanogaster. PLoS Genet, 11(5), e1005209. https://doi.org/10.1371/journal.pgen.1005209

Schlegel, P., Texada, M. J., Miroschnikow, A., Schoofs, A., Huckesfeld, S., Peters, M., Schneider-Mizell, C. M., Lacin, H., Li, F., Fetter, R. D., Truman, J. W., Cardona, A., & Pankratz, M. J. (2016). Synaptic transmission parallels neuromodulation in a central food-intake circuit. Elife, 5. https://doi.org/10.7554/eLife.16799

Scott, K., Brady Jr, R., Cravchik, A., Morozov, P., Rzhetsky, A., Zuker, C., & Axel, R. (2001). A chemosensory gene family encoding candidate gustatory and olfactory receptors in Drosophila. Cell, 104(5), 661–673.

Selcho, M., Muhlbauer, B., Hensgen, R., Shiga, S., Wegener, C., & Yasuyama, K. (2018). Anatomical characterization of PDF-tri neurons and peptidergic neurons associated with eclosion behavior in Drosophila. J Comp Neurol, 526(8), 1307–1328. https://doi.org/10.1002/cne.24408

Semaniuk, U. V., Gospodaryov, D. V., Feden’ko, K. M., Yurkevych, I. S., Vaiserman, A. M., Storey, K. B., Simpson, S. J., & Lushchak, O. (2018). Insulin-Like Peptides Regulate Feeding Preference and Metabolism in Drosophila. Front Physiol, 9, 1083. https://doi.org/10.3389/fphys.2018.01083

Shahid, S., Shi, Y., Yang, C., Li, J., Ali, M. Y., Smagghe, G., & Liu, T. X. (2021). CCHamide2-receptor regulates feeding behavior in the pea aphid, Acyrthosiphon pisum. Peptides, 143, 170596. https://doi.org/10.1016/j.peptides.2021.170596

Shiu, P. K., Sterne, G. R., Engert, S., Dickson, B. J., & Scott, K. (2022). Taste quality and hunger interactions in a feeding sensorimotor circuit. Elife, 11. https://doi.org/10.7554/eLife.79887

Siekhaus, D. E., & Fuller, R. S. (1999). A role for amontillado, the Drosophilahomolog of the neuropeptide precursor processing protease PC2, in triggering hatching behavior. Journal of Neuroscience, 19(16), 6942–6954.

Söderberg, J. A., Carlsson, M. A., & Nässel, D. R. (2012). Insulin-Producing Cells in the Drosophila Brain also Express Satiety-Inducing Cholecystokinin-Like Peptide, Drosulfakinin. Front Endocrinol (Lausanne), 3, 109. https://doi.org/10.3389/fendo.2012.00109

Sterne, G. R., Otsuna, H., Dickson, B. J., & Scott, K. (2021). Classification and genetic targeting of cell types in the primary taste and premotor center of the adult Drosophila brain. Elife, 10. https://doi.org/10.7554/eLife.71679

Sternson, S. M., Nicholas Betley, J., & Cao, Z. F. (2013). Neural circuits and motivational processes for hunger. Curr Opin Neurobiol, 23(3), 353–360. https://doi.org/10.1016/j.conb.2013.04.006

Sudhakar, S. R., Pathak, H., Rehman, N., Fernandes, J., Vishnu, S., & Varghese, J. (2020). Insulin signalling elicits hunger-induced feeding in Drosophila. Dev Biol, 459(2), 87–99. https://doi.org/10.1016/j.ydbio.2019.11.013

Talay, M., Richman, E. B., Snell, N. J., Hartmann, G. G., Fisher, J. D., Sorkac, A., Santoyo, J. F., Chou-Freed, C., Nair, N., Johnson, M., Szymanski, J. R., & Barnea, G. (2017). Transsynaptic Mapping of Second-Order Taste Neurons in Flies by trans-Tango. Neuron, 96(4), 783–795 e784. https://doi.org/10.1016/j.neuron.2017.10.011

Tatar, M., Kopelman, A., Epstein, D., Tu, M.-P., Yin, C.-M., & Garofalo, R. (2001). A mutant Drosophila insulin receptor homolog that extends life-span and impairs neuroendocrine function. Science, 292(5514), 107–110.

van den Pol, A. N. (2012). Neuropeptide transmission in brain circuits. Neuron, 76(1), 98–115. https://doi.org/10.1016/j.neuron.2012.09.014

Veenstra, J. A., Agricola, H. J., & Sellami, A. (2008). Regulatory peptides in fruit fly midgut. Cell Tissue Res, 334(3), 499–516. https://doi.org/10.1007/s00441-008-0708-3

Wang, P., Jia, Y., Liu, T., Jan, Y. N., & Zhang, W. (2020). Visceral Mechano-sensing Neurons Control Drosophila Feeding by Using Piezo as a Sensor. Neuron, 108(4), 640–650 e644. https://doi.org/10.1016/j.neuron.2020.08.017

Wang, Z., Singhvi, A., Kong, P., & Scott, K. (2004). Taste representations in the Drosophila brain. Cell, 117(7), 981–991. https://doi.org/10.1016/j.cell.2004.06.011

Watts, A. G., & Boyle, C. N. (2010). The functional architecture of dehydration-anorexia. Physiol Behav, 100(5), 472–477. https://doi.org/10.1016/j.physbeh.2010.04.010

Williams, M. J., Akram, M., Barkauskaite, D., Patil, S., Kotsidou, E., Kheder, S., Vitale, G., Filaferro, M., Blemings, S. W., Maestri, G., Hazim, N., Vergoni, A. V., & Schioth, H. B. (2020). CCAP regulates feeding behavior via the NPF pathway in Drosophila adults. Proc Natl Acad Sci U S A, 117(13), 7401–7408. https://doi.org/10.1073/pnas.1914037117

Wu, M., Nern, A., Williamson, W. R., Morimoto, M. M., Reiser, M. B., Card, G. M., & Rubin, G. M. (2016). Visual projection neurons in the Drosophila lobula link feature detection to distinct behavioral programs. Elife, 5. https://doi.org/10.7554/eLife.21022

Yapici, N., Cohn, R., Schusterreiter, C., Ruta, V., & Vosshall, L. B. (2016). A Taste Circuit that Regulates Ingestion by Integrating Food and Hunger Signals. Cell, 165(3), 715–729. https://doi.org/10.1016/j.cell.2016.02.061

Yeom, E., Shin, H., Yoo, W., Jun, E., Kim, S., Hong, S. H., Kwon, D. W., Ryu, T. H., Suh, J. M., Kim, S. C., Lee, K. S., & Yu, K. (2021). Tumour-derived Dilp8/INSL3 induces cancer anorexia by regulating feeding neuropeptides via Lgr3/8 in the brain. Nat Cell Biol, 23(2), 172-183. https://doi.org/10.1038/s41556-020-00628-z

Yoshinari, Y., Kosakamoto, H., Kamiyama, T., Hoshino, R., Matsuoka, R., Kondo, S., Tanimoto, H., Nakamura, A., Obata, F., & Niwa, R. (2021). The sugar-responsive enteroendocrine neuropeptide F regulates lipid metabolism through glucagon-like and insulin-like hormones in Drosophila melanogaster. Nat Commun, 12(1), 4818. https://doi.org/10.1038/s41467-021-25146-w

Yu, H. H., Awasaki, T., Schroeder, M. D., Long, F., Yang, J. S., He, Y., Ding, P., Kao, J. C., Wu, G. Y., Peng, H., Myers, G., & Lee, T. (2013). Clonal development and organization of the adult Drosophila central brain. Curr Biol, 23(8), 633–643. https://doi.org/10.1016/j.cub.2013.02.057

Zhai, R. G., & Bellen, H. J. (2004). The architecture of the active zone in the presynaptic nerve terminal. Physiology, 19(5), 262–270.

Zhang, F., Mak, S. O. K., Liu, Y., Ke, Y., Rao, F., Yung, W. H., Zhang, L., & Chow, B. K. C. (2022). Secretin receptor deletion in the subfornical organ attenuates the activation of excitatory neurons under dehydration. Curr Biol, 32(22), 4832–4841 e4835. https://doi.org/10.1016/j.cub.2022.09.037

Zhang, L., Guo, X., & Zhang, W. (2022). Nutrients and pheromones promote insulin release to inhibit courtship drive. Science Advances, 8(10), eabl6121.

Zheng, Z., Lauritzen, J. S., Perlman, E., Robinson, C. G., Nichols, M., Milkie, D., Torrens, O., Price, J., Fisher, C. B., Sharifi, N., Calle-Schuler, S. A., Kmecova, L., Ali, I. J., Karsh, B., Trautman, E. T., Bogovic, J. A., Hanslovsky, P., Jefferis, G., Kazhdan, M., . . . Bock, D. D.. (2018). A Complete Electron Microscopy Volume of the Brain of Adult Drosophila melanogaster. Cell, 174(3), 730–743 e722. https://doi.org/10.1016/j.cell.2018.06.019

Zimmerman, C. A., Leib, D. E., & Knight, Z. A. (2017). Neural circuits underlying thirst and fluid homeostasis. Nature Reviews Neuroscience, 18(8), 459–469.

